# Systematic evaluation of parameters for genome-scale metabolic models of cultured mammalian cells

**DOI:** 10.1101/2020.06.24.169938

**Authors:** Song-Min Schinn, Carly Morrison, Wei Wei, Lin Zhang, Nathan E. Lewis

## Abstract

Genome-scale metabolic models describe cellular metabolism with mechanistic detail. Given their high complexity, such models need to be parameterized correctly to yield accurate predictions and avoid overfitting. Effective parameterization has been well-studied for microbial models, but it remains unclear for higher eukaryotes, including mammalian cells. To address this, we enumerated model parameters that describe key features of cultured mammalian cells – including cellular composition, bioprocess performance metrics, mammalian-specific pathways, and biological assumptions behind model formulation approaches. We tested these parameters by building thousands of metabolic models and evaluating their ability to predict the growth rates of a panel of phenotypically diverse Chinese Hamster Ovary cell clones. We found the following considerations to be most critical for accurate parameterization: (1) cells limit metabolic activity to maintain homeostasis, (2) cell morphology and viability change dynamically during a growth curve, and (3) cellular biomass has a particular macromolecular composition. Depending on parameterization, models predicted different metabolic phenotypes, including contrasting mechanisms of nutrient utilization and energy generation, leading to varying accuracies of growth rate predictions. Notably, accurate parameter values broadly agreed with experimental measurements. These insights will guide future investigations of mammalian metabolism.

## Introduction

Cultured mammalian cells are prominent expression systems for large-scale biotherapeutics manufacturing. However, conventional bioprocess engineering has largely been empirical due to lacking knowledge of cell biology. Recent advances in high dimensional data and computational methods are improving and accelerating industrial cell line and bioprocess development^1–3^. In particular, metabolism has been a key target in cell line development^4–8^, given its role in meeting the anabolic and bioenergetic needs of cell growth and protein production^9,10^. Correspondingly, there has been increased interest to investigate the metabolism of mammalian cells, such as Chinese hamster ovary (CHO) cells, using genome-scale metabolic network models^11–18^. These models contextualize large-scale biological data with curated biochemical knowledge, and have been used with a wide array of *in silico* methods^19,20^ to probe the molecular basis of metabolism^21,22^, disease^23–25^, and human-microbiome interactions^26,27^. More recently, metabolic network models are being applied to industrial bioprocess development^28–30^ – e.g. predict metabolic phenotype^31,13,32^, identify metabolic bottlenecks^16^, and optimize media formulation^15,33^.

A general challenge of genome-scale metabolic network modeling is to identify relevant insights from an underdetermined solution space^34^, all the while avoiding overfitting. That is, these models typically consist of thousands of metabolites, and reactions that far outnumber experimental measurements available as boundary conditions, often leading to an underdetermined system that is vulnerable to overfitting. To address this challenge, the following established steps can be taken: (1) create a context-specific sub-model consisting of the most relevant metabolic reactions for a given context by interpreting gene expression data^17,35,36^, (2) input experimental data as boundary conditions, and (3) hypothesize likely metabolic flux values by applying optimality principles to the model reaction network^37–39^. Each of these steps invite the modeler to make assumptions about cellular features – e.g. cellular makeup^40,41^, the relevance of specific pathways^42,43^ and ‘cellular objectives’^38,39,44–46^ – that improve the model’s solution space reflect biology. Here, we define the model implementation of these assumptions as ‘model parameters.’ In the last two decades, such model parameters have been extensively explored and refined for model microbes such as *Escherichia coli*^47,48^, *Saccharomyces cerevisae*^49,50^ and others^51,52^. However, parameters for higher eukaryotes remain under-characterized, despite notable recent advances^53–55^.

Mammalian cells are distinct from microbes in their metabolic pathways, auxotrophic requirements, sub-compartmentalization and regulation^56^, and therefore require parameterization that capture such differences. A few studies have investigated mammalian-specific parameters describing cell biomass^57^, novel metabolic pathways^12^ and cellular objectives^58,14^. These parameters, however, have not been rigorously evaluated against wide-ranging conditions; meanwhile, other features of mammalian metabolism and growth remain unexplored altogether. To address this, we systematically investigate a panel of parameters for describing diverse cultured mammalian cell metabolic phenotypes, with a focus on CHO cells. Our work identifies key parameters for predicting metabolic phenotypes, challenges conventional parameterization approaches, and guides future modeling efforts for mammalian metabolism.

## Results

### A cell line-specific model was constructed for phenotypically diverse CHO clones

Cultured mammalian cells, such as CHO cells, consume media nutrients – e.g. glucose, amino acids and cofactors – to produce biomass, recombinant proteins and byproducts. Depending on genotype and environmental conditions, CHO cells vary in their metabolic rates and pathway usage, resulting in high metabolic heterogeneity. Here, we have examined a variety of CHO metabolic phenotypes to identify general modeling principles and avoid overfitting. Specifically, we investigated 10 CHO clones that differed in recombinant antibody, bioreactor conditions and gene knockout treatment (see Methods). Consequently, the cells exhibited varying patterns of nutrient consumption and byproduct secretion (Fig. 1A), ultimately leading to divergent growth and productivity phenotypes (Fig. 1B; Fig. S1). We also observed the 10 clones between culture days 4 to 11 (Fig. 1C) for temporal variation in metabolism^59,60^.

**Figure 1:**
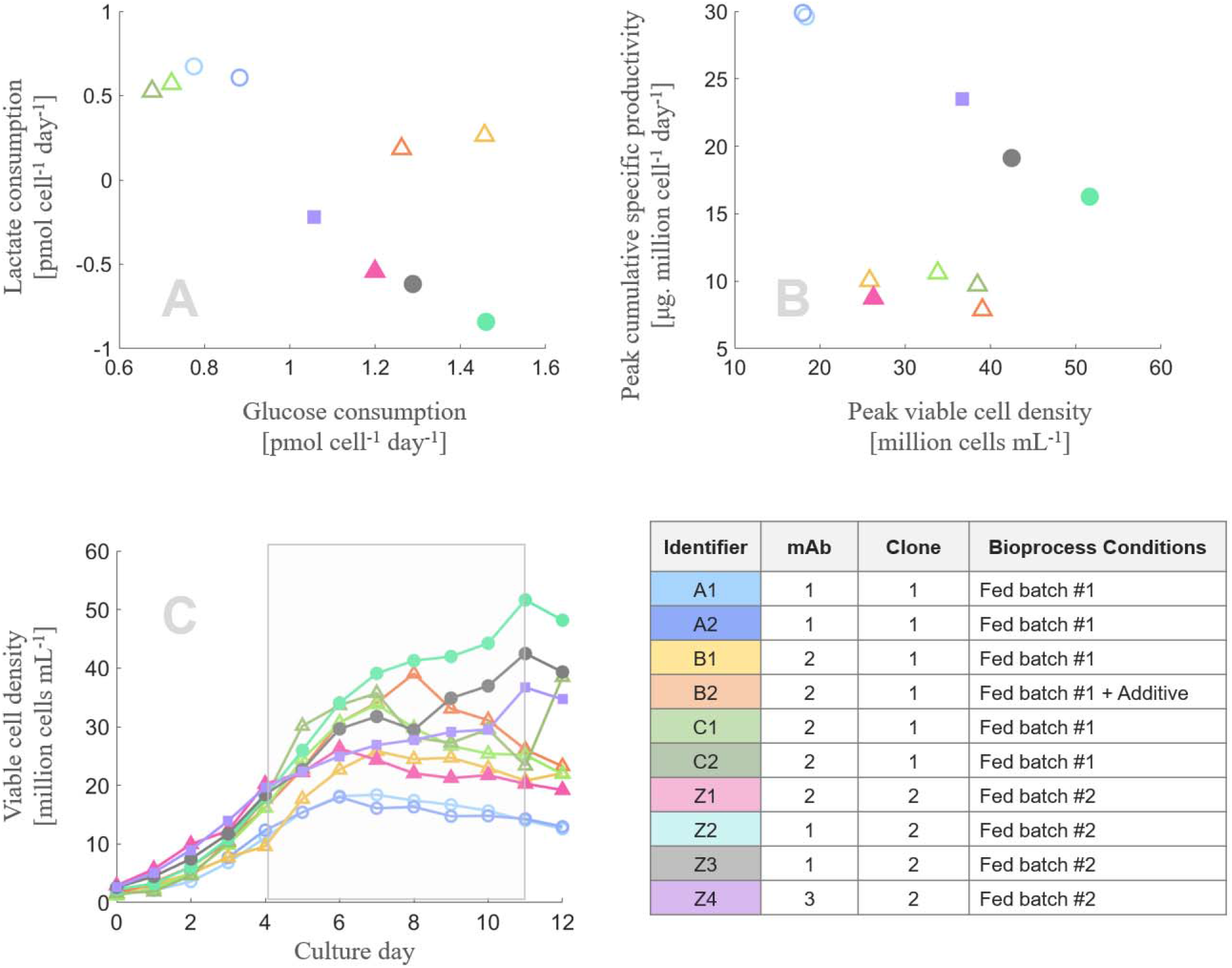
10 CHO clones with diverse metabolic phenotypes were studied. The examined clones expressed different monoclonal antibodies (1: ○, 2: △, 3: □) and were subjected to different bioprocess conditions (Fed batch #1: empty; Fed batch #: filled). Due to these differences, the cells exhibited diverse metabolic phenotypes: (a) the cells consumed key nutrients such as glucose at varying amounts, and variously consumed or secreted lactate, (b) resulting in distinct growth and productivity performances. (c) These clones were observed between culture days 4 and 11, during which the cells traversed exponential and stationary phases (highlighted).

Broadly, we saw several distinct types of metabolisms and nutrient utilization efficiencies. Some cells displayed high glucose consumption and high proliferation, suggesting an efficient and fast-moving metabolism (e.g. clones Z2, Z3). Other cells showed low glucose consumption and low proliferation but high protein production, suggesting an efficient but attenuated metabolism (e.g. clones A1, A2). Still others displayed high glucose consumption but low proliferation and low protein production, suggesting a severely inefficient metabolism under cellular stress.

We hypothesized that these diverse CHO metabolisms could be described by a well-parameterized metabolic network model. To test this, we constructed a cell line-specific metabolic network model by modifying a previously published genome-scale model of CHO metabolism^11^. Briefly, this was done by the following steps: (1) transcriptomics data were analyzed to quantify the relative activity of 210 metabolic tasks to hypothesize a list of active metabolic genes^36,61,62^, (2) these active genes and their associated model reactions were refined into a fully functional model via the mCADRE algorithm^63^, and (3) the subsequent draft model was manually curated.

### Mammalian-specific parameters embedded biological assumptions onto model

To enable the curated model of describing diverse CHO metabolisms, we identified salient features of mammalian cell cultures and formulated them as model parameters (Table 1; see Supplementary Document for detailed and technical discussions on their computational implementation, including annotated code). First, we considered cellular biomass’ macromolecular makeup and dry weight (Table 1, parameters 1-2). These parameters are necessary for the model to describe proliferation via the biomass production reaction^37^, and to convert experimental measurements (*per cell* units) into model-compatible values (*per cellular dry weight* units). In other words, these parameters are critical to bounding the model solutions to experimental data. While the importance of these parameters are well-established previously^57,41,64,40^, here we compare them with other novel parameters in the context of mammalian cell biology.

**Table 1:**
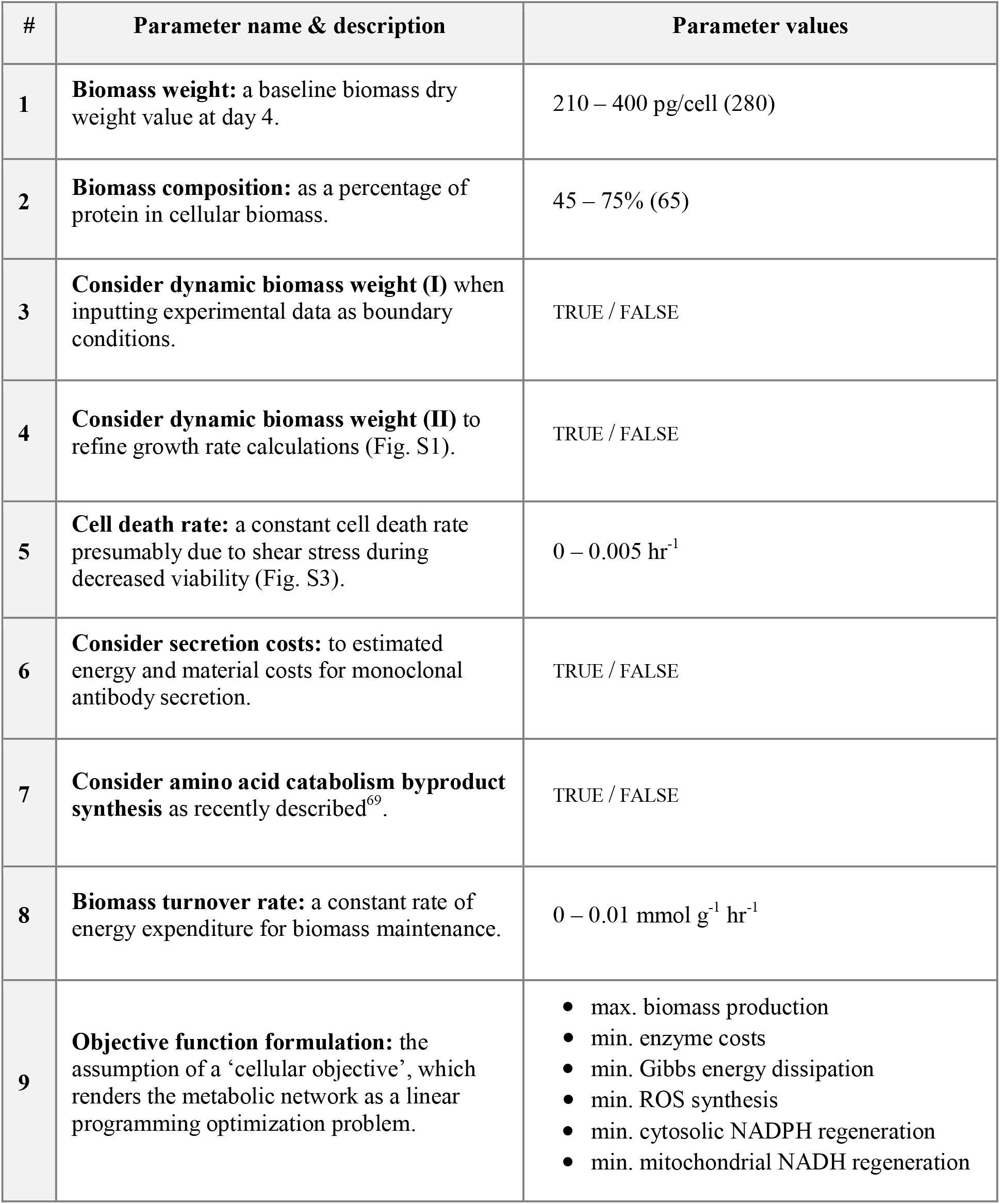
Parameters varied in this study. the table provides names and descriptions of the parameters, along with range of values explored in our analysis; mean experimental values of the presented clones are given in parentheses when applicable.

| # | Parameter name & description | Parameter values |
| --- | --- | --- |
| 1 | **Biomass weight:** a baseline biomass dry weight value at day 4. | 210 – 400 pg/cell (280) |
| 2 | **Biomass composition:** as a percentage of protein in cellular biomass. | 45 – 75% (65) |
| 3 | **Consider dynamic biomass weight (I)** when inputting experimental data as boundary conditions. | TRUE / FALSE |
| 4 | **Consider dynamic biomass weight (II)** to refine growth rate calculations (Fig. S1). | TRUE / FALSE |
| 5 | **Cell death rate:** a constant cell death rate presumably due to shear stress during decreased viability (Fig. S3). | 0 – 0.005 hr<sup>-1</sup> |
| 6 | **Consider secretion costs:** to estimated energy and material costs for monoclonal antibody secretion. | TRUE / FALSE |
| 7 | **Consider amino acid catabolism byproduct synthesis** as recently described<sup>69</sup>. | TRUE / FALSE |
| 8 | **Biomass turnover rate:** a constant rate of energy expenditure for biomass maintenance. | 0 – 0.01 mmol g<sup>-1</sup> hr<sup>-1</sup> |
| 9 | **Objective function formulation:** the assumption of a ‘cellular objective’, which renders the metabolic network as a linear programming optimization problem. | • max. biomass production<br>• min. enzyme costs<br>• min. Gibbs energy dissipation<br>• min. ROS synthesis<br>• min. cytosolic NADPH regeneration<br>• min. mitochondrial NADH regeneration |

Second, we considered that cellular dry weight change during fed-batch culture, indicating internal shifts in metabolism and productivity^65^. Indeed, in the presented experiments, cellular mass increased by as much as 40-70% throughout the culture period (Fig. S2). Up to now, modeling efforts have treated this feature as static, even for multi-week time-course studies. A dynamic cellular dry weight could facilitate more accurate experiment-to-model unit conversions. It could also refine calculations between proliferation and biomass production. For example, it would enable the model to consider that CHO cells may still produce significant amounts of biomass during stationary phase when they proliferate more slowly but grow in size (see also Fig. S2C). The impact of these considerations on model results is unclear.

We also observed cell loss despite high viability (>99%) during the transition between exponential growth phase and stationary phase (Fig. S4), presumably due to shear stress-induced apoptosis^66,67^. The cell loss was primarily observed on clones grown in Fed Batch #1 (Fig. 1, Fig S4A). Accounting for these lost cells could improve calculations of total biomass produced during later stages of cell culture. We explore these several bioprocess-related phenomena (Table 1, parameters 3-5) as novel model parameters.

Third, we improved descriptions of mammalian metabolic pathways which are not found in microbes (Table 1, parameters 6-8). For example, we included in the model an intricate multi-organelle secretory pathway^68^ to improved estimations of metabolic expenditures towards producing complex recombinant proteins (Table S3). Similarly, we also considered metabolic pathways for producing various byproducts from amino acid catabolism^69^ (Table S4), and cellular death and turnover^70^.

Fourth, we reevaluated the biological premises behind applying the optimality principle to the metabolic network model (Table 1, parameter 9). This mathematical operation assumes that cellular metabolism operates at some optimal condition and is crucial to constraining the metabolic network model. As an example, proliferative bacteria have often been described to *maximize biomass production* for rapid growth, which is computationally formulated as a biomass objective function^37^. This objective function presumes that biomass production is principally limited by nutrient availability. Meanwhile, a variety of alternative objective functions have also been explored^38,39^ – e.g. optimal energy generation per substrate^71^, minimized redox potential^58,72,73^, efficient use of enzyme capacity^74,75^ and streamlined amino acid transport^14^. Compared to microbes, mammalian cells are larger, more complex and wired for multicellularity, all of which increase the costs of homeostatic maintenance and limit proliferation. Therefore, we compare the conventional *maximize biomass production* function against alternative objective functions describing mammalian metabolism as chiefly limited by various homeostasis requirements – e.g. thermal^76^, proteomic^77,78^, reactive oxygen species or redox homeostasis^79^. These alternative objective functions were realized by imposing ‘penalties’ on model reactions (see Methods; Table S5, 6).

### Experimental measurements help produce robust model parameters

We investigated how the above-described model parameters impacted model prediction of metabolism. The nine parameters were randomly permuted in value to within-literature-reported ranges thousands of times. A resulting set of specified parameter values is called here as a ‘parameter setting’. In total, 4000 distinct parameter settings were generated and used to bound the solution space. We tested these parameter settings against experimental data for 10 clones between culture days 4 to 11, resulting in 320,000 model predictions (Fig. 2a; Table S7). Each of these predictions yielded a model-predicted growth rate that could be compared to experimental measured growth rates (Fig. 2b). For ease of interpretation, prediction errors were preprocessed to yield an ‘accuracy’ metric (Fig. 2c). Specifically, the prediction errors were transformed by a negative-log function and normalized so that the average error metric for all clones and timepoints would lie between 0 and 1. For reference, when growth rates were predicted to within 5% of experimental measurements for all 80 points, the accuracy metric equaled 0.8.

**Figure 2:**
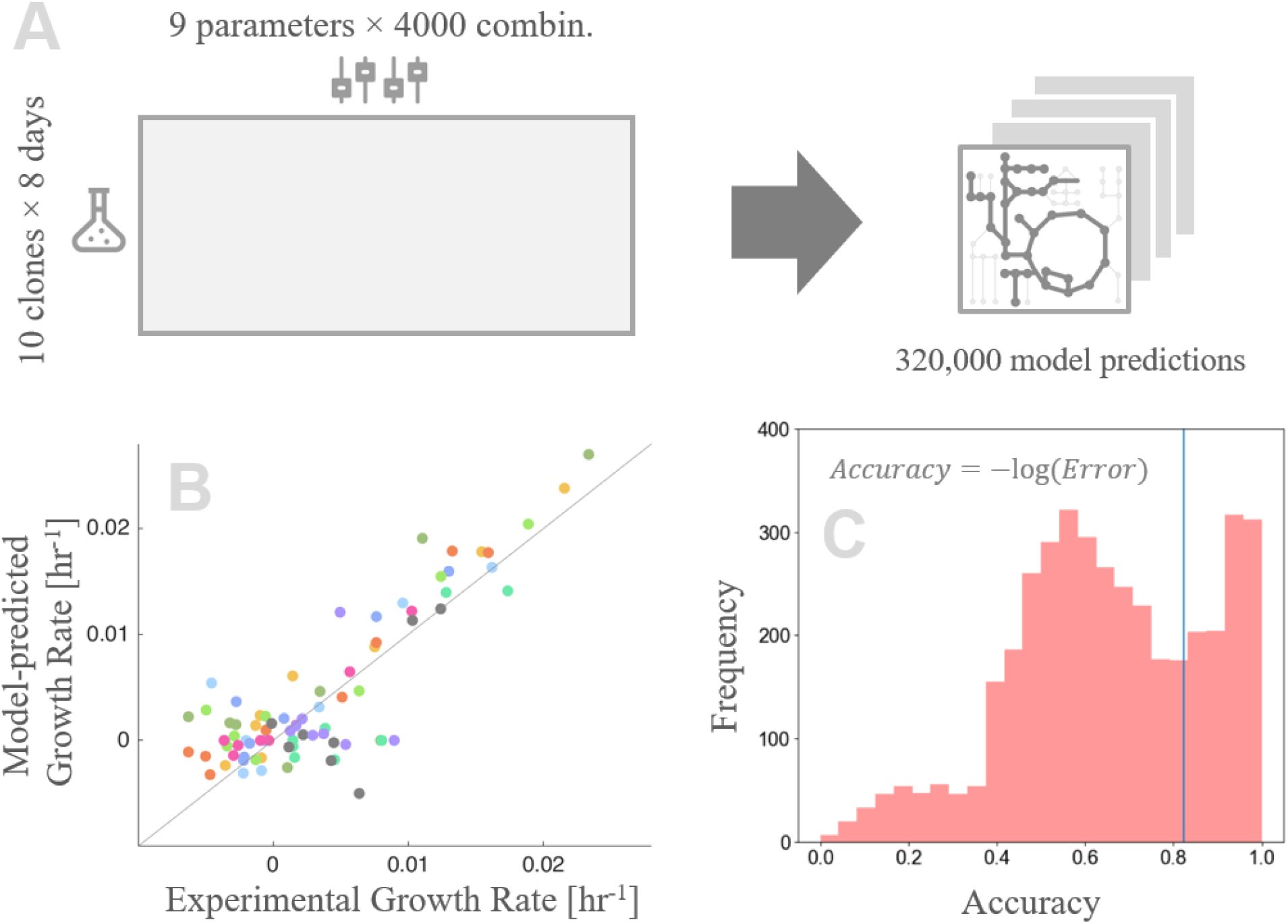
Analysis of model parameters, workflow scheme & results. (a) 4000 parameter settings were evaluated against 80 datapoints to produce thousands of model predictions. (b) Here, we provide an example model prediction of growth rate for 80 points. These particular predictions were made by assuming parameter values equal to experimental measurements – that is, a baseline biomass dry weight of 280 picograms/cell at day 3, and 65% protein composition of biomass; biomass weight was also varied according to time-course cell diameter measurements. Biomass production was assumed to be limited by enzyme capacity. (c) Analysis results were examined by their ‘accuracy’, which was calculated and normalized from the negative-log of mean prediction residuals. Predictions assuming experimental values, featured in panel (b), resulted in an average accuracy of 0.82 (blue line).

Overall, the various parameter settings resulted in a wide range of prediction accuracies (Fig. S4). About one-eighth of the parameter settings robustly described at least eight of the ten clones with high accuracy (accuracy > 0.8). Half of the parameter settings predicted only two or fewer clones with high accuracy, indicating that the models were highly overfit or poorly parameterized. The remaining one-third of the parameter settings predicted a moderate range of clones with high accuracy, suggesting various degrees of overfitting. Overfit parameters of one clone tended to also describe biologically similar clones better, recapitulating known differences in cell line lineage and bioprocess conditions (Fig. S5).

Experimental data helped produce robust models. A model parameterized according to experimental measurements (Table 1, parenthesized) yielded an average accuracy of 0.82 across all clones and timepoints (Fig. 2b; 2c, blue line). Likewise, parameter settings with values close to experimental measurements (within ±15%) resulted in comparable accuracies (0.83 ± 0.11). Only a minority of parameter settings (2%) significantly deviated from experimental values yet managed to perform comparably. The value of experimental measurements was especially apparent for the *cell death rate* parameter describing late-stage cell death observed only in some clones. The *cell death rate* parameter helped the model recapitulate observed cross-clonal variation (Fig. S6).

### Selection of objective function has greatest influence on model prediction

We estimated the importance of the parameters for prediction accuracy using a linear regression model (Fig. 3). Of the nine parameters, *objective function formulation* had an outsized influence on prediction accuracy. Specifically, the *maximize biomass production* function was correlated strongly with poor predictions. In contrast, the *minimize cytosolic NADPH regeneration* function was well correlated with accurate predictions. The following parameters also improved model predictions: *cell death rate*, *biomass composition* and *consideration of dynamic dry weights to calculate growth rate* (Fig. S8). Interestingly, the time-course of biomass weight was more important than the baseline weight value itself. The remaining parameters had negligible impact on model accuracy (effect size < 0.01; see Fig. S7).

**Figure 3:**
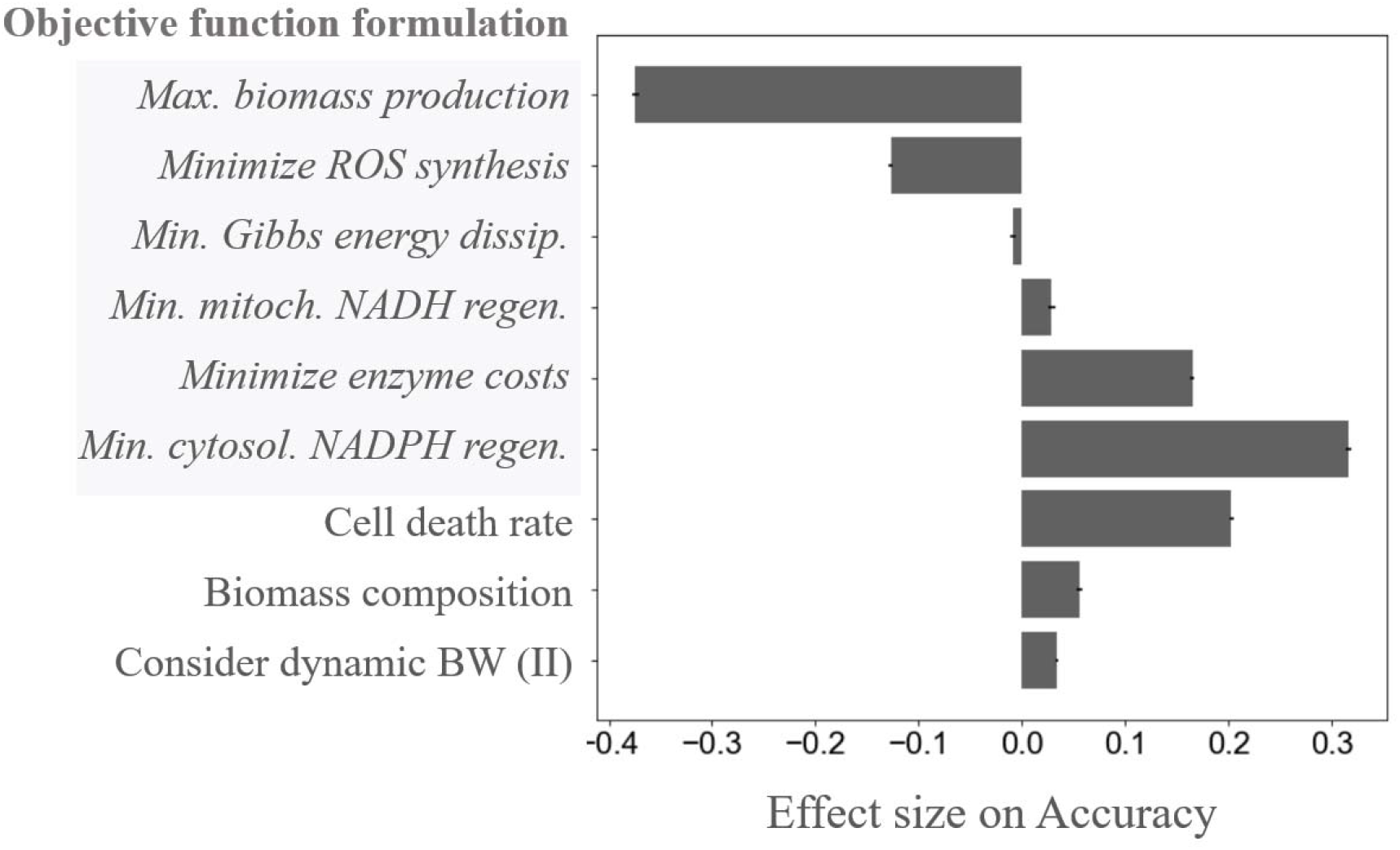
Key model parameters impact model accuracy. A regression model estimated the importance of parameters by their effect size. *Objective function formulation* was treated as six independent binary parameters representing the various formulations (italicized and highlighted in grey). Several of these formulations were highly correlated to prediction accuracy. Other important parameters were *cell death rate*, *biomass composition* and *consideration of dynamic biomass weight* (BW)*for growth rate calculations*. The remaining parameters had negligible effects, which are detailed elsewhere (Fig. S6)

The *objective function formulations* provided detailed and mechanistic hypotheses of the intracellular metabolic activities underlying the growth rate predictions (Fig. 4), as revealed by flux sampling analysis (see Methods). The *maximize biomass production* objective function predicted an efficient and highly active metabolism, which utilized almost all consumed substrates towards generating energy and biomass. As a result, the model consistently overestimated growth rates across clones and timepoints (Fig. S9). Overestimations were particularly pronounced for inefficient metabolisms, such as clones B1 and B2. In contrast, several homeostasis-related objective functions predicted less efficient metabolisms that partially discard consumed nutrients via various shunting mechanisms. This assumption, supported by carbon balance estimations (Fig. S10), was key to predicting CHO metabolisms with differing nutrient utilization efficiencies in a well-rounded manner.

**Figure 4:**
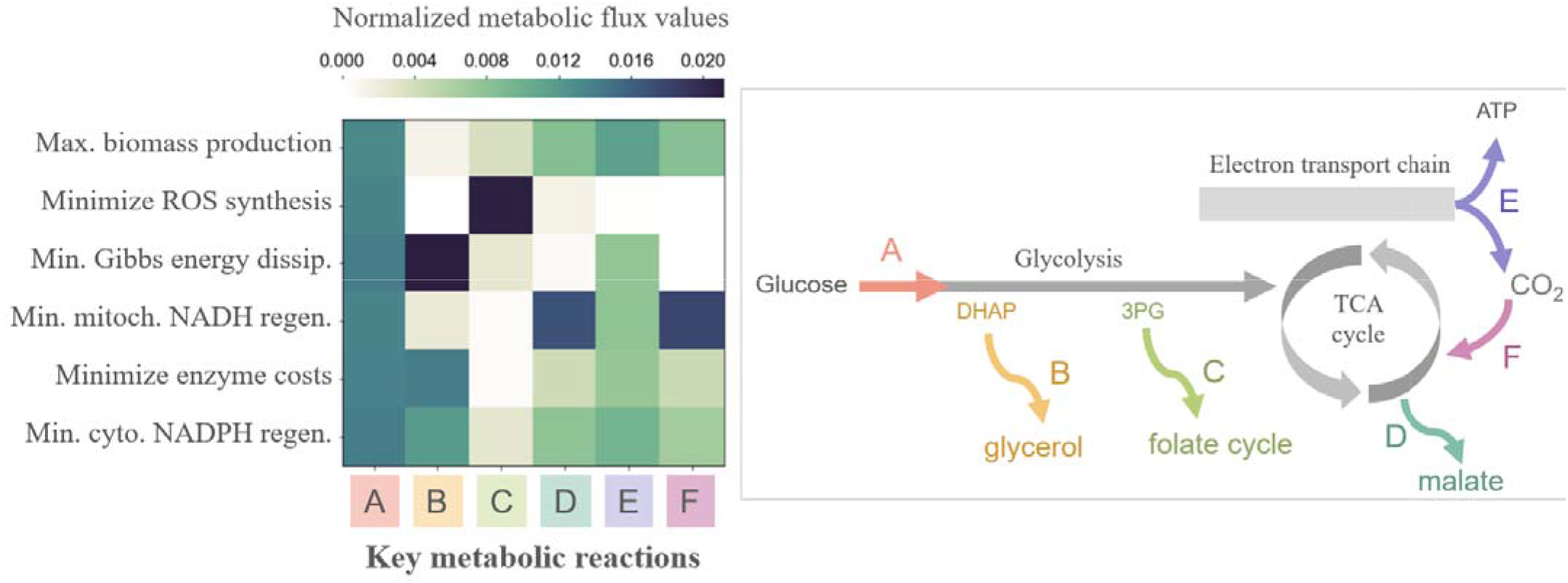
Objective functions vary in hypothesized metabolic activities underlying the predicted growth rates. Objective functions helped determine the metabolic activities of key reactions in the central carbon metabolism. The heatmap displays the reactions’ fractional contribution to total metabolic flux. These reactions include the first step of glycolysis catalyzed by hexokinase 1 (*A*); carbon shunting variously towards glycerol (*B*), folate cycle intermediates (*C*), malate secretion (*D*); mitochondrial respiration (*E*); anaplerosis of respiratory carbon dioxide via bicarbonate and pyruvate carboxylase (*F*). The flowchart on the right contextualizes the reactions within the central carbon metabolism.

Specifically, the homeostasis-related objective functions hypothesized two different carbon shunting mechanisms: (1) secretion of glycerol, which is converted from the glycolytic intermediate dihydroxyacetone phosphate via glycerol-3-phosphate dehydrogenase and glycerol-3-phosphate phosphatase (Fig. 3, *B*), and (2) secretion of malate, an intermediate of the citric acid cycle and malate-aspartate shuttle (Fig. 3, *D*). The objective functions *minimize enzyme costs*, *minimize Gibbs energy dissipation* and *minimize cytosolic NADPH regeneration* hypothesized shunting via glycerol, with varied fluxes through glycolysis, glycerol synthesis, mitochondrial respiration and pyruvate carboxylate. The objective function *minimize mitochondrial NADH regeneration* hypothesized shunting via malate. This loss of flux in the citric acid cycle was then counteracted by converting carbon dioxide from respiration to bicarbonate, to eventually oxaloacetate via pyruvate carboxylate^80^ (Fig. 3, *F*). The secretions of both glycerol and malate have been reported previously in CHO fed-batch systems^81,82,10^, although only glycerol was observed in the present study to accumulate in the spent medium (Fig. S11).

Lastly, the *minimize ROS synthesis* objective function hypothesized negligible activities in glycolysis and citric acid cycle, deviating from known CHO biology^83^. Instead, most consumed glucose was diverted to the folate cycle via phosphoserine transamination to produce energy and byproduct formic acid (Fig. 3, *C*), resulting in an improbable metabolic phenotype. Additional comparison of objective functions and their influence on metabolic configurations are also provided (Fig. S12).

## Discussion

Parameters help constrain metabolic networks sufficiently and accurately by embedding knowledge and data onto models. We compared thousands of distinct parameter settings for their ability to describe CHO clones with wide-ranging phenotypes. The resulting analyses lead to important insights for parameterizing metabolic network models of cultured mammalian cells. Specifically, this study (1) confirms that experimental data improve model parameterization, (2) identifies relevant parameters for mammalian metabolic models, most prominently the *objective function formulation*, and (3) challenges the popular use of the *maximize biomass production* objective function for modeling exponentially growing mammalian cells and explores promising alternatives.

Our analysis of model parameters agreed with experimental observations in several notable ways. First, experimentally measured parameter values resulted in robust and accurate model predictions, confirming the value of experimental measurements in parameterization. Second, parameter settings that described one clone well also tended to describe similar clones well, recapitulating differences in cell line lineages and glucose-feeding strategies. Lastly, parameter values of *cell death rate* recapitulated experimental observations at the clone level. These broad agreements demonstrate that well-parameterized metabolic network models can describe diverse metabolic states.

Our analysis identified parameters that strongly affected model precision. Of all the investigated parameters, the *objective function formulation* had a predominant influence on model-predicted growth rates and their underlying metabolic activities. *Biomass composition* was also reconfirmed as a key parameter, agreeing with previous findings in microbial models^40,64,41^. Lastly, novel parameters accounting for time-course changes in cell size and viability improved model predictions substantially. This emphasizes the relevance of dynamic and bioprocess-specific features of mammalian cell cultures for parameterization.

Notably, the widely-used *maximize biomass production* objective function performed poorly in describing mammalian metabolism. For microbial metabolism, this formulation is supported by theoretical and experimental evidence^84^, and has been widely predictive for many experimental conditions and different models^39,73^. Accordingly, several recent studies using mammalian metabolic models (e.g., CHO cells, cancer cells, stem cells, immune cells) adopted this assumption. In the present study, however, *maximize biomass production* consistently overestimated growth rates and intracellular metabolic activity. Alternatively, five other presented formulations assumed that mammalian cells limit biomass production to maintain various homeostatic conditions. These homeostasis-limited assumptions performed markedly better than *maximize biomass production* by hypothesizing a restrained CHO metabolism. Objective functions considering cytosolic redox homeostasis and enzyme capacity predicted metabolic configurations that agreed especially well with observed growth rates and exometabolomic measurements. Future studies can rigorously refine and validate these formulations by comparing predicted intracellular flux distributions with experimental flux measurements, following the well-established footsteps of *Escherichia coli* models^38,39^.

## Conclusion

Proper parameter selection is essential for metabolic network models to provide accurate descriptions of cellular metabolism. Above all, the objective function parameter is highly influential in predicting metabolic and bioprocess performance phenotype. For this purpose, mammalian cells may be described as limiting their metabolic activities to maintain homeostasis. Various objective functions predicted different growth rates and metabolic pathway usage, leaving room for further validation and refinements. In addition, parameters describing biomass and time-course metabolic shifts also improved model predictions, especially when set to experimentally measured values. These results will guide future efforts to develop and improve models for mammalian cell metabolism, with applications ranging from biotherapeutic production^18,85^ to unraveling the metabolic basis of diverse diseases^86–88^.

## Methods

### Cell culture experiments

Two production fed batch processes were used, Fed batch 1 and Fed batch 2. Both fed batch processes used chemically defined media and feeds over the 12-day cell culture. Fed batch 1 used a glucose restricted fed batch process called HiPDOG^65^. Glucose concentration is kept low during the initial phase of the process, Day 2-7, through intermittent addition of feed medium containing glucose at the high end of pH dead-band and then glucose was maintained above 1.5 g/L thereafter. These conditions help restrict lactate production in fed batch cultures without compromising the proliferative capability of cells. In Fed batch 2 a conventional cell culture process was used where glucose was maintained above 1.5 g/L throughout the process.

For both process conditions, bioreactor vessels were inoculated at 2 × 10^6^ viable cells/mL. The following bioprocess characteristics were quantified daily using a NOVA Flex BioProfile Analyzer (Nova Biomedical, Waltham, MA): viable cell density, average live cell diameter and concentrations of glucose, lactate, glutamate, and glutamine. Viable cell density (VCD) data was converted to growth rates by following equation to be compared to model-predicted growth rates.

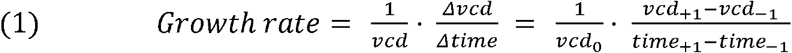

In addition, cell cultures were sampled on specific days (Day 0, 3, 5, 7, 10 and 12) for cell pellets and supernatant for transcriptomic (RNA-Seq), metabolomics analyses, and titer measurements. Titers were analyzed using a protein A HPLC (model 1100 HPLC, Agilent Technologies, Inc., Santa Clara, CA, protein A column model 2-1001-00, Applied Biosystems, Foster City, CA). The RNA libraries were mapped to the CHO genome^89,90^ using STAR aligner^91^ and processed to quantify gene expression counts with HTSeq-count^92^.

Flash-frozen cell pellets (10E6 cells) and supernatant (1 mL) were collected from bioreactor runs for clones A1 and A2 for each sampling day. Similarly, cell pellet and supernatant samples were collected from bioreactor runs for clones B1 and B2 for days 7, 10 and 12. Collected samples were sent to Metabolon (Metabolon Inc, Morrisville, NC) for metabolomics analyses. Proteins were removed by methanol precipitation and the metabolites were recovered by vigorous shaking and centrifugation. The extracted samples were run for reverse-phase Ultrahigh Performance Liquid Chromatography-Tandem Mass Spectroscopy with negative ion mode ESI. Raw data was extracted, peak-identified and processed for quality control using Metabolon’s hardware and software. The raw ion count data was normalized against the extracted proteins quantified using a Bradford assay.

### Biomass measurement experiments

During cell culturing of clones Z1-Z4, 20 × 10^6^ cells were sampled during days 0, 3, and 7. At day 7, an additional 1 mL of culture was sampled. The samples were sub-divided and stored at −80 °C for the following measurements.

Cellular dry weights were measured by weighing dehydrated cells, as previously described^93^. Briefly, for each sample, an aluminum weigh boat was dried at 70 °C for 48 hours and pre-weighed after being cooled down to room temperature. Stored samples (5 × 10^6^ cells) were thawed in ice, centrifuged, washed with PBS and centrifuged again. The cell pellets were resuspended in deionized water and transferred to the pre-weighed aluminum boats and dried for 48 hours at 70° C. Then, the samples were cooled back to room temperature and weighed in triplicates. The weighed mass was divided by cell count to calculate cellular dry weights.

Cellular protein contents were quantified by the Bicinchoninic Acid assay using a commercial kit (Pierce™ BCA Protein Assay Kit; Thermo Fisher, Waltham, MA). Stored samples (1 × 10^6^ cells) were thawed in ice, centrifuged, and washed with PBS. Assay standards were prepared as instructed for the range of 0 – 2000 μg/mL protein. The cells were disrupted by a mixture of cell lysis agent (CelLytic™ M; Sigma Aldrich, St. Louis, MO) and protease inhibitor cocktail (Sigma Aldrich, St. Louis, MO). A negative control was prepared from the mixture without cell samples. The standards, samples and control were treated with assay reagents, incubated and measured following kit instructions. The resulting standard curve was confirmed to be linear (R^2^ = 0.988). Measured absorbance values were converted to protein concentrations following this standard curve and considering background absorbance described by the negative control. Protein content in biomass was calculated to be 64.9% (±6.8) for days 3 and 7, agreeing well with previously published measurements^11,57^.

The lipid contents of the cells were extracted using the Blight and Dyer protocol^94^. Briefly, for each bioreactor run, 1 mL stored culture samples were centrifuged and re-suspended in water. Chloroform and methanol were added sequentially in 1:2 ratio and mixed well by vortex. Then, chloroform and water are sequentially added in 1:1 ratio and mixed well in between by vortex. The resulting mixture was then centrifuged at 4000 RPM for 15 minutes at 20 °C, resulting in phase separation. The upper phase and cell debris were carefully discarded. The lower chloroform phase containing the lipids was carefully transferred to pre-weighed glass beakers. Chloroform evaporated in a fume hood for 24 hours, after which the lipid samples were weighed. The measured weights were divided by cell count and measured cellular dry weight, yielding a cellular lipid content of 21.62% (±2.46), agreeing well with previously published measurements.

### Constraint-based modeling and analysis

Flux balance analyses were conducted using the COBRA Toolbox 2.0^95^ and the Gurobi solver version 8.0.0 (Gurobi Optimization; Beaverton, Oregon) in MATLAB R2018b (MathWorks; Natick, Massachusetts, USA).

### Constructing a context-specific model

A community-consensus genome-scale model^11^ was modified to produce a single cell line-specific model for the cell lines from Pfizer by qualitatively interpreting transcriptomics data for clone A1, following benchmarking results36,61,62. Specifically, transcript abundance data for each gene was pre-processed by equation 2 to produce a binary gene score metric (ON: >5, OFF: 0). The equation’s threshold values were calculated from the mean transcript abundance across all samples. Genes with transcript abundances of top and bottom 25 percentile were binarized as ON and OFF, respectively. The binary gene scores were converted to reaction scores by considering multimeric or isozyme relationships as described by the model’s gene-protein-relationship matrix. These binary reaction scores were used as input data for model extraction by the mCADRE algorithm^63^.

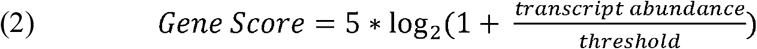

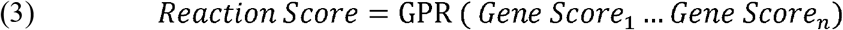

The resulting draft model was then manually curated to be consistent with experimental data and established cell biology. Given the defined media of Pfizer’s bioprocess, we allowed extracellular transport only for metabolites with experimental basis – e.g. glucose, lactate, proteinogenic amino acids, oxygen, carbon dioxide etc. We confirmed amino acid essentialities and CHO-specific auxotrophies for arginine, cysteine and proline. We also ensured the inclusion of synthesis pathways of non-essential amino acids and degradation pathways for all amino acids. We confirmed the model utilized glucose via central carbon metabolism reactions, and removed unlikely reactions – e.g. the methylglyoxal pathway or cross-membrane transport of metabolites not found in media formulation. We ensured cytosolic and mitochondrial compartmentalization of redox cofactors such as NAD/H and NADP/H. The final model consisted of 2375 irreversible reactions, 587 metabolites and 1043 genes (Table S1, S2).

### Model implementation of parameters

Here we detail how the nine parameters were implemented in the model. The workflow and decision-making described here are also visualized as a flowchart (Fig. S13). The parameter value of *biomass weight* (i.e. cellular dry weight) was used to convert the units of experimental measurements from [pg·cell^−1^·day^−1^] to [mmol·g_DW_^−1^·hr^−1^]. When *considering dynamic dry weight (I)* for data input, the dry weight values were estimated from cell diameter measurements by assuming a constant cellular density and geometry^96^. When *considering dynamic dry weight (II)* for growth rate calculations, the following equation was used.

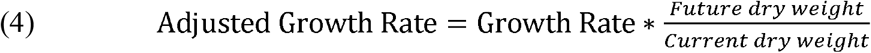

The *biomass composition* parameter was adjusted by altering the stoichiometric formulation of the biomass production reaction. Specifically, macromolecular protein content and lipid content were inversely varied, while assuming content of nucleotides and carbohydrates were stable. This was because protein and lipid contents were observed to vary the most^57^. Molecular makeup of amino acids and specific lipid molecules was unchanged from the genome-scale model, which was based on approximations from hybridoma cells. This was because variations in molecular composition did not affect model predictions. The *cell death rate* was implemented by the following equation for timepoints between days 7-11, in accordance to experimental observations.

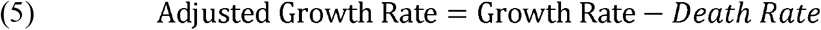

*Consideration of secretion costs* was implemented by joining a model of the CHO secretory pathway^68^ onto our metabolic network model. The secretory pathway model included about 100 reactions for folding, transport, post-translational modification and related activities in the endoplasmic reticulum and Golgi body (Table S3). We approximated the structure, folding requirements and glycosylation needs of Pfizer’s monoclonal antibodies by that of Rituximab. *Biomass turnover rate* was implemented a single ATP hydrolysis reaction.

*Consider byproduct synthesis* was implemented for the following byproduct molecules: glycerol, formic acid, 2-hydroxybutyurate, isovalerate, acetic acid, hydroxyphenylpyruvate, hydroxyphenyl-lactate, phenyl-lactate, homocysteine, indole 3-lactate, citrate, and malate. These notably include products of amino acid catabolism as well as citric acid cycle intermediates.

Glycerol was assumed to be synthesized from dihydroxyacetone phosphate via glycerol-3-phosphate dehydrogenase and glycerol-3-phosphate phosphatase. Formic acid was assumed to be synthesized in the folate cycle during tetrahydrofolate synthesis from 10-formyltetrahydrofolate. The synthesis of isovalerate, 2-hydroxybutyurate, acetic acid, hydroxyphenylpyruvate, hydroxyphenyl-lactate, phenyl-lactate, homocysteine, and indole 3-lactate were formulated as catabolic byproducts, as previously published^69^. The model already included synthesis reactions for homocysteine, acetic acid, citrate and malate. Synthesis and transport reactions were enabled for all byproducts (Table S4).

### Formulation of the objective function

The most widely used objective function is the maximization of biomass production, which formulates the model as a linear programming optimization problem around the biomass production reaction^97^. Alternatively, we formulated the objective function as a two-step linear programming optimization to reflect cellular limitations to biomass production. First, we enumerated possible limitations to biomass production, as discussed above. Second, we annotated model reactions for ‘penalty’ metrics describing such limitations (Table S5, S6). We derived penalty values for Gibbs dissipation^76^ and enzyme costs^98^ from previous works. Third, the model was bounded by experimentally measured nutrient consumption rates (Table S8). Fourth, for a penalty of choice, we calculated the minimal amount of penalty to process the consumed nutrients. This calculated value was then used as an additional boundary condition. Fifth, given all these bounds, we calculated the maximum possible amount of biomass production. This predicted value, then, describes the maximum anabolic activity given nutrient consumption behavior and theoretical cellular limitations.

This presented workflow builds upon previous work that minimizes network flux^99^ and parsimonious flux balance analysis^78^ and the ‘max biomass per unit flux’ objective^38^ with important differences. Previous methods implicitly assumed that biomass production was gated by substrate availability and therefore first maximized biomass production and then minimized total unit flux as a penalty metric. Here, we assume cellular limitations restrict growth rather than substrate availability, and therefore inverse the order of optimization. Also, while previous methods penalized reactions uniformly, we apply weighted penalties following biological assumptions. For example, reactions that are catalyzed by complex multimers are penalized correspondingly more severely. Like previous methods, this workflow requires the model to be irreversible.

### Estimation of parameter impact on model accuracy

4000 parameter settings were generated by randomly varying parameters to within a range based on literature or experimental measurements (Table 1, Table S7). For each setting, parameter values were implemented to yield a flux balance analysis prediction. The prediction used as input data the following experimental measurements: specific productivity and the consumption rates of glucose, lactate and 20 proteinogenic amino acids (Table S8). The prediction was repeated 80 times by using experimental data from 10 clones across 8 days. The resulting 80 model-predicted growth rates were compared to their respective experimental values to yield prediction errors (Table S7). To facilitate human interpretation, the prediction errors were first transformed by the negative log function. Then, the transformed values were normalized so that mean metric values for all 80 predictions laid between 0 and 1. The resulting metric was called ‘accuracy’.

A regression analysis between accuracy and parameter values were performed, using Python 3.7.3 in the Jupyter Notebook environment. For this analysis, parameter values were normalized to be between 0 and 1 by referencing minimum and maximum possible values (Table 1). A linear regression analysis was performed according to equation 6, where B_*i*_ and X_*i*_ represent respectively the effect size and normalized parameter values of parameter *i*. The resulting effect size of parameters and their p-values were used to identify parameter relevance.

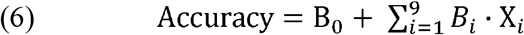

### Flux sampling analysis

We computed distributions of likely fluxes for each reaction in the model by stochastically sampling 5000 points within the solution space via the Markov chain Monte Carlo sampling via artificially centered hit-and-run algorithm, as described previously^100^. First, the metabolic network model was parameterized according to experimentally observed values. Then, experimental measurements for all clones on day 4 were used to constrain model reactions for biomass production, monoclonal antibody secretion and consumption of glucose, lactate and proteinogenic amino acids. A set of non-uniform ‘points’ or flux values was generated within the feasible flux space. Each point was subsequently moved randomly, while remaining within the feasible flux space. To do this, a random direction was first chosen. Second, the limit for how far the point can travel in the randomly-chosen direction was calculated. Lastly, a new random point on this line was selected. This process was iterated until the set of points approached a uniform sample of the solution space. Thereafter, the sampled fluxes were normalized by total model flux^101^. The normalized values were analyzed to explore in detail the impact of objective functions on model predictions. These analyses were done using Python 3.7.3 in the Jupyter Notebook environment.

## Supporting information

Annotated Pseudocode

