## Supplementary Figures for "Systematic evaluation of parameters for genome-scale metabolic models of cultured mammalian cells"

**Contents**

Figure S1: Overview of phenotypic diversity among investigated clones

Figure S2: Dynamic cellular morphology in CHO bioprocess

Figure S3: Cell death rate and ‘disappearing’ cells

Figure S4: Parameter settings predict growth rates with varying accuracies

Figure S5: Parameter settings recapitulate known phenotypic differences

Figure S6: Parameter settings recapitulates clonal variations observed in cell death rate

Figure S7: Parameters by effects size on model accuracy

Figure S8: Parameter effects on model accuracy

Figure S9: Maximize biomass production consistently overestimates growth rates

Figure S10: Estimated carbon balance suggests inefficient metabolism

Figure S11: Metabolomics data suggest glycerol secretion

Figure S12: Objective functions produce diverse intracellular flux distributions

Figure S13: Flowchart of model parameterization

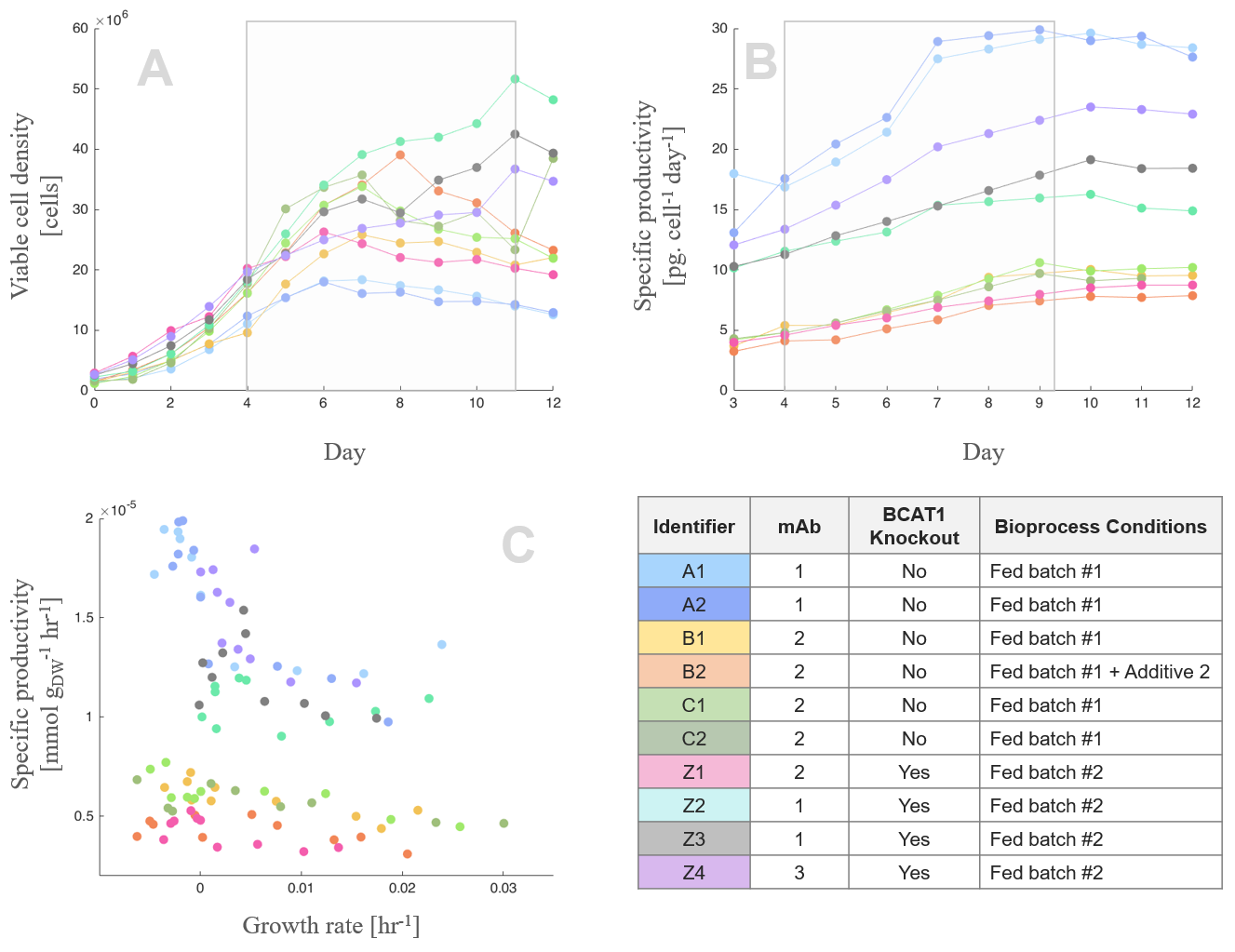

**Figure S1: Overview of phenotypic diversity among investigated clones**

Cell culture experiments of 10 clones – Ten industrial producing Chinese Hamster Ovary cells were investigated. The clones varied in their parental cell line, recombinant antibody and bioprocess conditions (inset table, lower right). Consequently, the clones exhibited differing growths and productivities across culture periods (a, b). The diversity of surveyed metabolic states is shown (c). The highly proliferative phenotype is found in the bottom right, while high-productivity phenotype is found in the top left. Less optimal phenotypes displaying lower growth and productivity is found in bottom left.

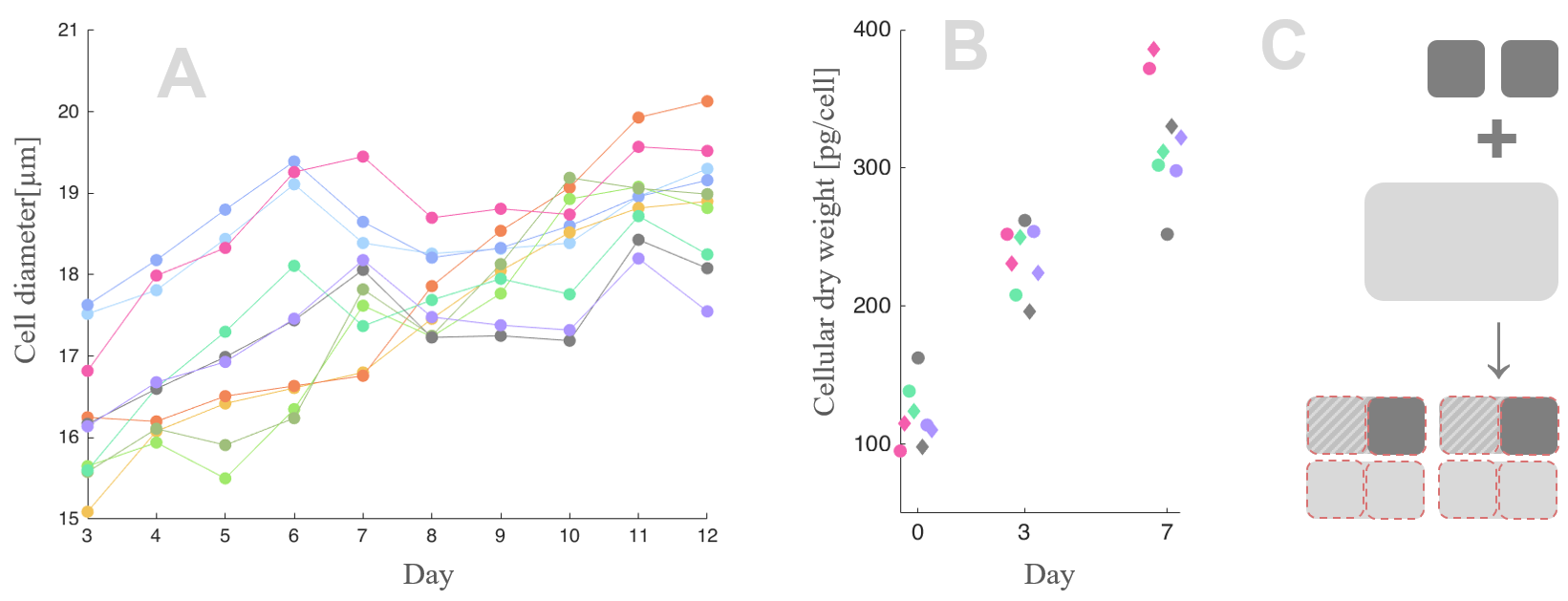

**Figure S2: Dynamic cellular morphology in CHO bioprocess**

(a) We observed significant changes in cell diameters throughout culture period in all ten clones, suggesting that cellular dry weight was changing as well. Overall, observed cellular diameter increased 10-20% across the entire culture period. (b) We confirmed time-course changes in dry weight for four clones from day 0 to 7. (c) A mock example is provided to illustrate the potential impact of changing dry weight on growth calculations. In this example, two cells (top, dark grey squares) produce biomass (middle, light grey) to both expand and proliferate (bottom). The produced biomass is used to double cell size (shaded) and also create two new larger cells (bottom, light grey rectangles). If a constant cell size is assumed, the produced biomass would correspond to six new cells (bottom, red dotted outlines), which overestimates proliferation rate by two-fold.

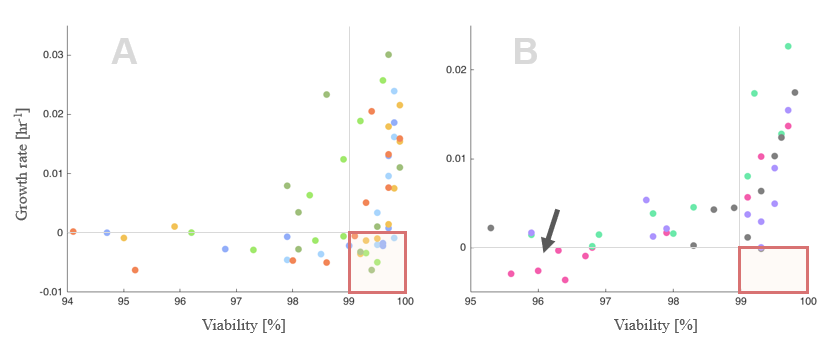

**Figure S3: Cell death rate and ‘disappearing’ cells**

The growth rate and viability are plotted for 10 clones. Early on in the culture, cells exhibit fast growth and high viability (upper right). Throughout the culture, cells slow growth and reduce in viability, and therefore ‘traverse’ gradually downward and leftward. During the transition between growth and stationary phase, (a) cell lines A, B and C exhibited decreasing cell density despite high viability (bottom right, red box). We hypothesize this to be due to a small portion of unviable cells being degraded into cell debris from shear stress. (b) Cell line Z did not exhibit such phenomenon (empty red box). Only clone Z1 (pink markers; arrow) exhibited decreasing cell density during days 7-11. Apoptosis due to shear stress has been observed to be cell line- or process-specific previously^1^. Clonal variation in this cell death rate phenotype was recapitulated during parameter sensitivity analysis (Fig. S6)

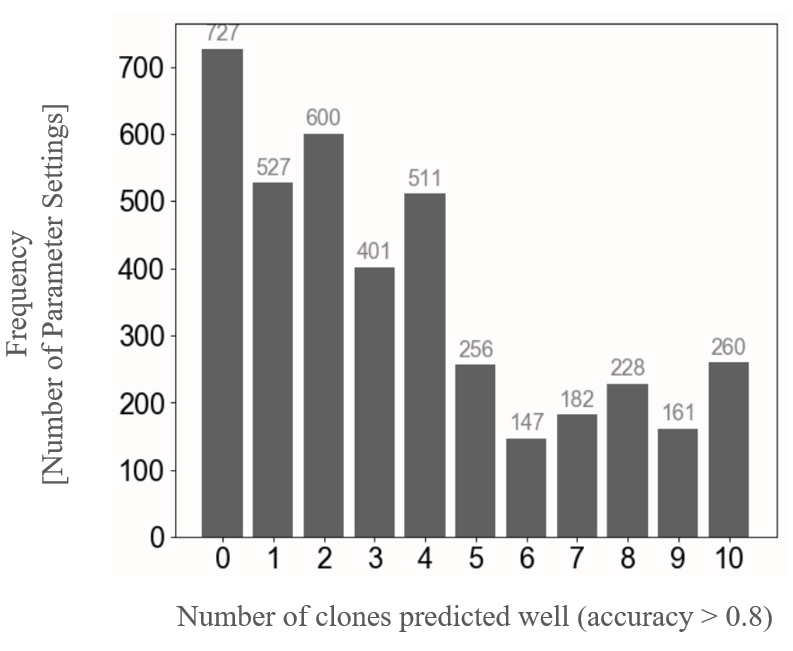

**Figure S4: Parameter settings predict growth rates with varying accuracies**

For each of the 4000 parameter settings, we evaluated for how many of the 10 clones it could produce growth rate predictions with high accuracy. High accuracy was defined by the 75^th^ percentile value of all accuracies (accuracy^75%tile^ = 0.7949).

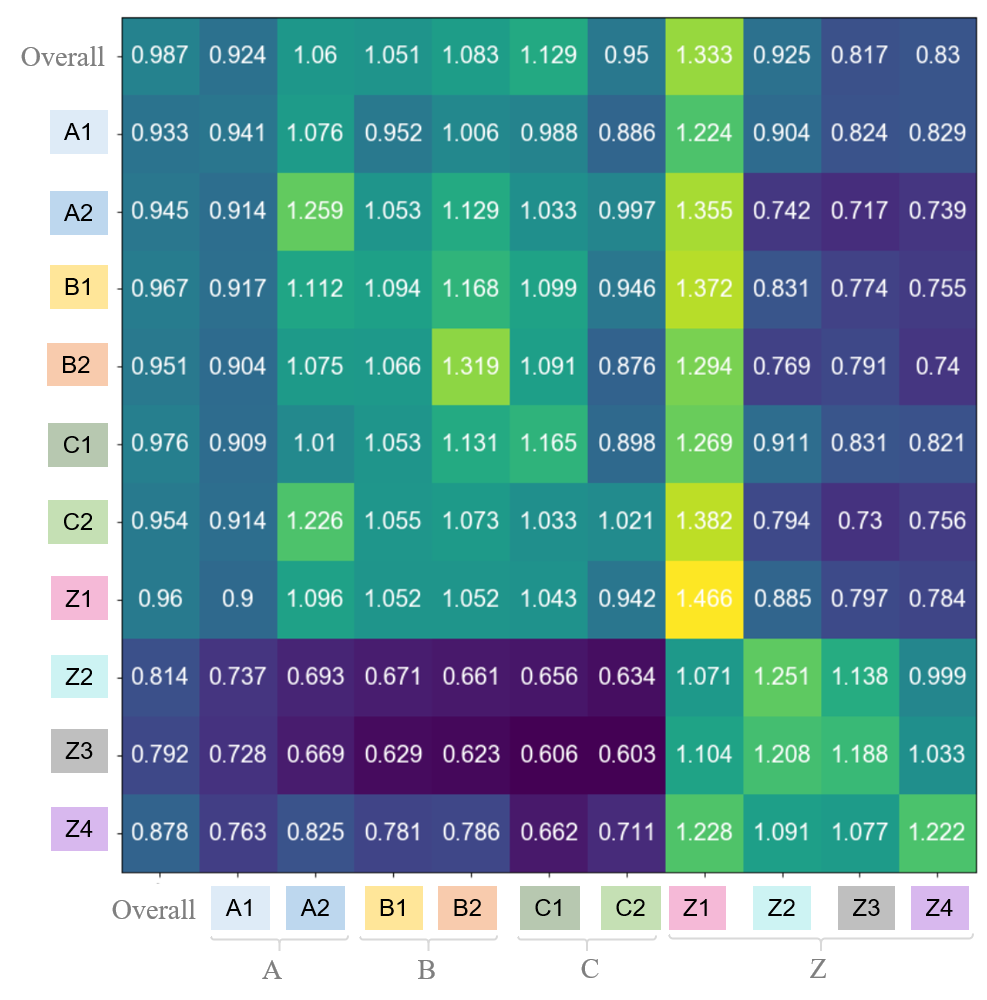

**Figure S5: Parameter settings recapitulate known phenotypic differences**

For each clone, parameter settings with highest accuracies were identified (row). Then, these top-performing parameter settings were evaluated across other clones for accuracy (column). For example, optimal parameter settings for clone Z2 (mean accuracy = 1.251) result in accuracies of 0.634 and 1.071 for clones C2 and Z1, respectively. Stark differences in parameter performances were observed across cell lines A-C and Z, which differ in cell line lineage and glucose feeding regimen. Note that prediction accuracy for a given clone could be greater than 1, since all accuracy values was normalized based on average performance across all clones and timepoints.

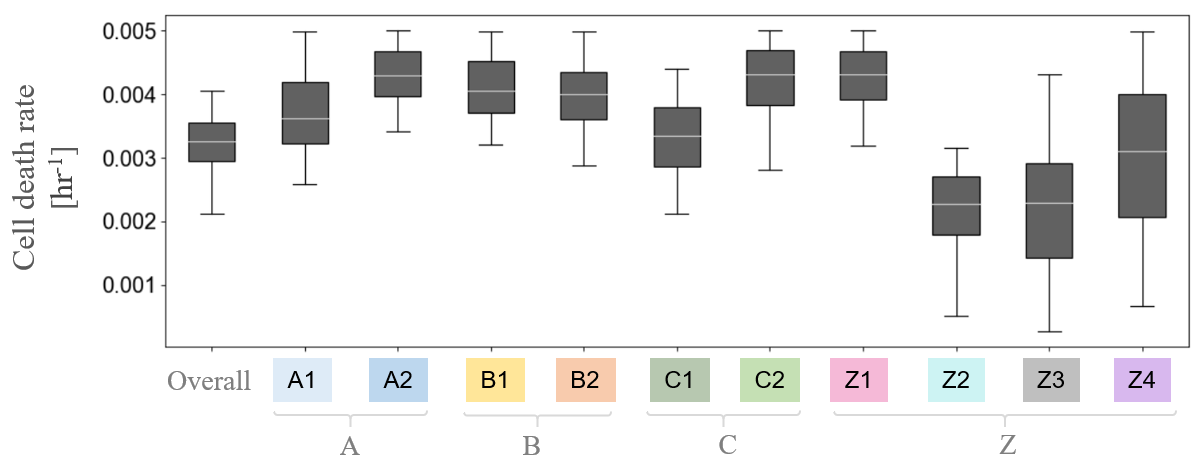

**Figure S6: Parameter settings recapitulates clonal variations observed in cell death rate**

For each clone, we identified 100 parameter settings with highest accuracies and noted their corresponding *cell death rate* values. Relatively high cell death rates (>0.003 hr^-1^) resulted in higher accuracies for cell lines A, B and C, as well as clone Z1. In contrast, lower cell death rates performed better for clone Z2. Lastly, a variety of cell death rates performed well for clones Z3 and Z4, suggesting that cell death rate was less influential in determining accuracy for these predictions. This overall trend corresponds remarkably well with experimental findings, where cell death rate was observed for cell lines A, B, C and clone Z1, but not for other clones of cell line Z (Fig. S3).

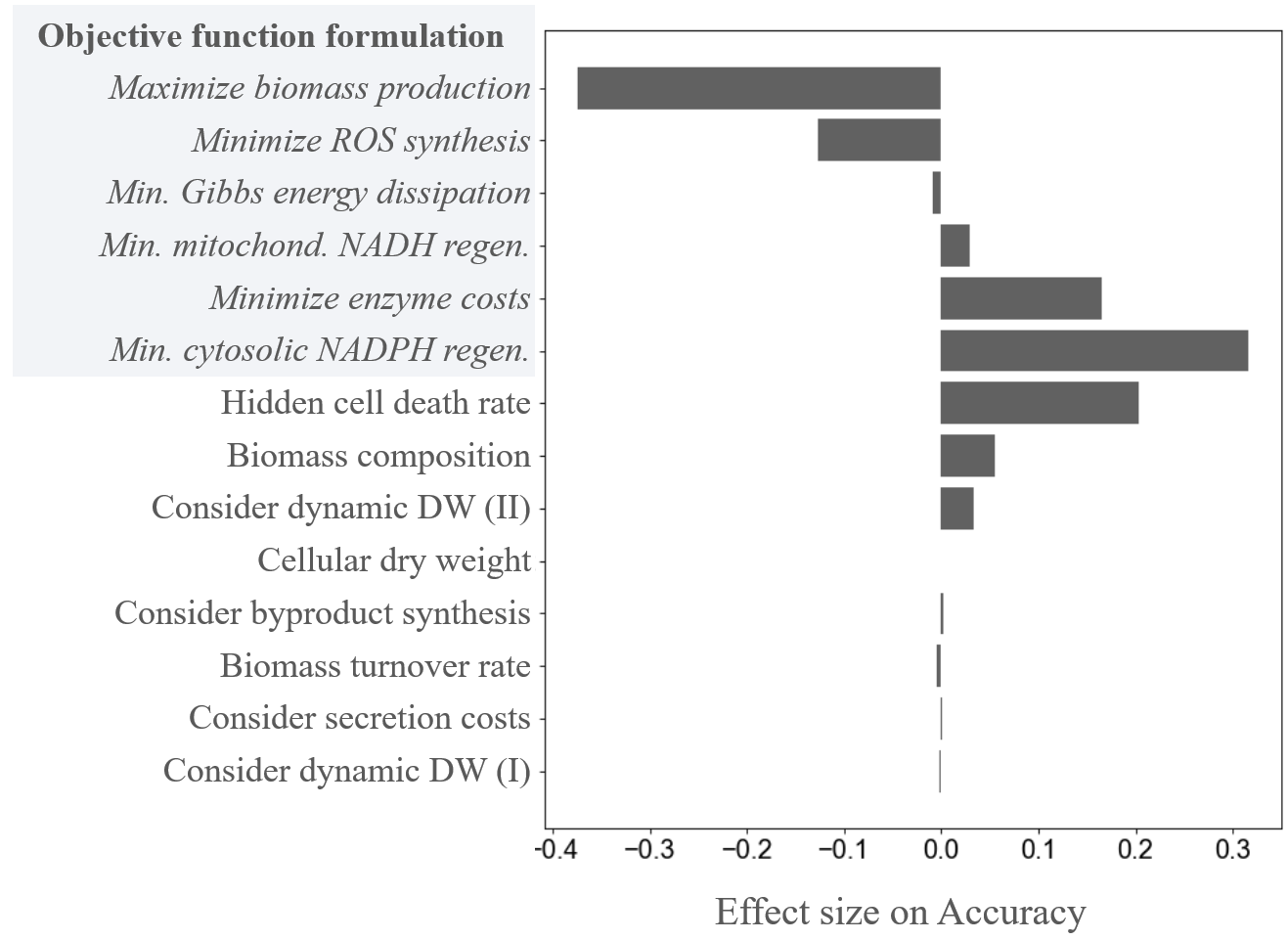

**Figure S7: Parameters by effects size on model accuracy**

Regression analysis estimates the importance of each parameter by their effect size. The formulation of the objective function was by far the most influential parameter (shaded in grey).

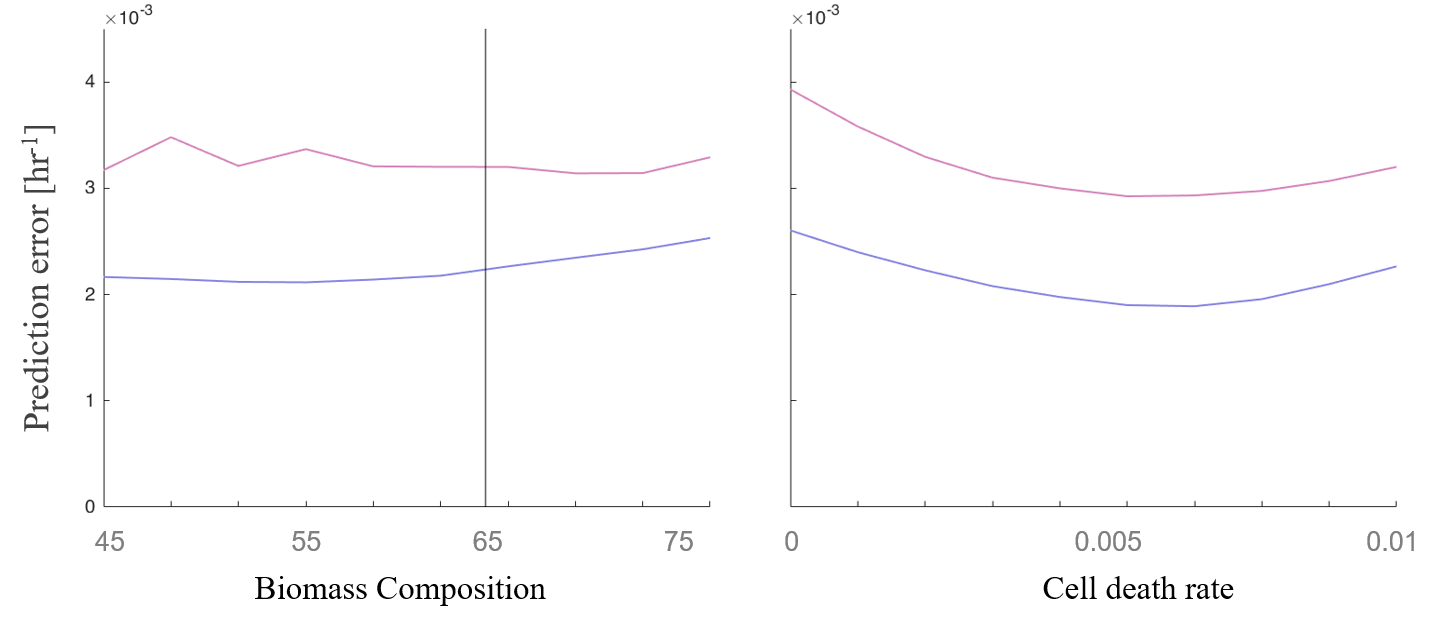

**Figure S8: Parameter effects on model accuracy**

The effects of the parameters *biomass composition* and *cell death rate* are explored. Biomass composition is expressed as a percentage of protein in cellular biomass. Mean prediction errors from exponential phase and stationary phase timepoints are shown in purple and magenta, respectively. Experimentally measured protein percentage is marked by a vertical line (65%).

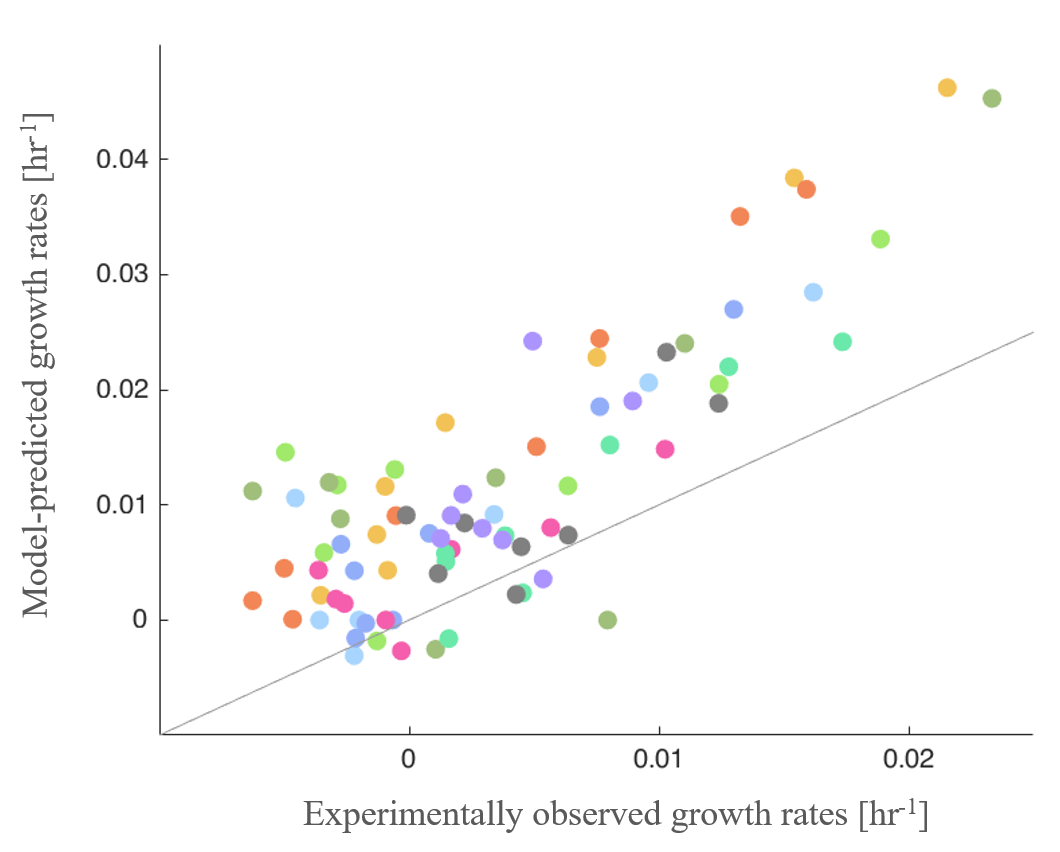

**Figure S9: *Maximize biomass production* consistently overestimates growth rates**

Growth rates for 10 clones across 8 culture days were predicted using the *maximize biomass production* objective function. Other parameter values were set to experimental measurements – i.e. a *biomass weight* of 280 picogram/cell at day 3, and *biomass composition* at 65% protein. Further, biomass weight was varied dynamically based on cellular diameter measurements. The resulting predictions consistently overestimate growth rates.

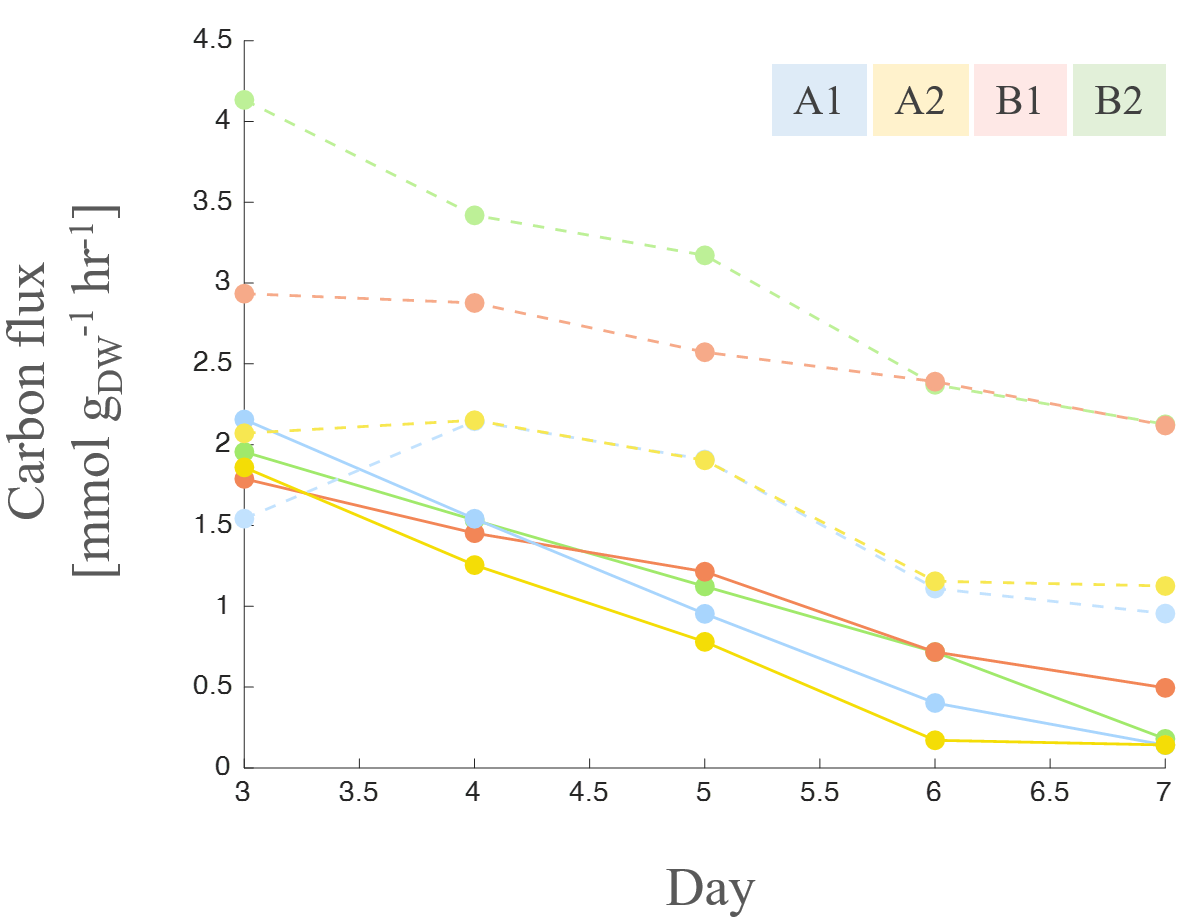

**Figure S10: Estimated carbon balance suggests inefficient metabolism**

We estimated carbon flux involved in consumption (dotted) and anabolic activities (solid).

We estimated the total carbon flux associated with the consumption of glucose, lactate and amino acids (dotted) and with anabolic activities of biomass production and transgenic protein synthesis (solid). Comparison of such carbon ‘income’ and ‘expenditures’ for cell lines A and B show that Chinese Hamster Ovary cells can uptake high amounts of carbon which they do not use for biomass production. Consumed carbon flux was estimated from key metabolites’ stoichiometric makeup and consumption rates. Anabolic carbon flux was estimated by assuming a cellular stoichiometric makeup of CH_1.61_O_0.56_N_0.16_, found in budding yeast^2^.

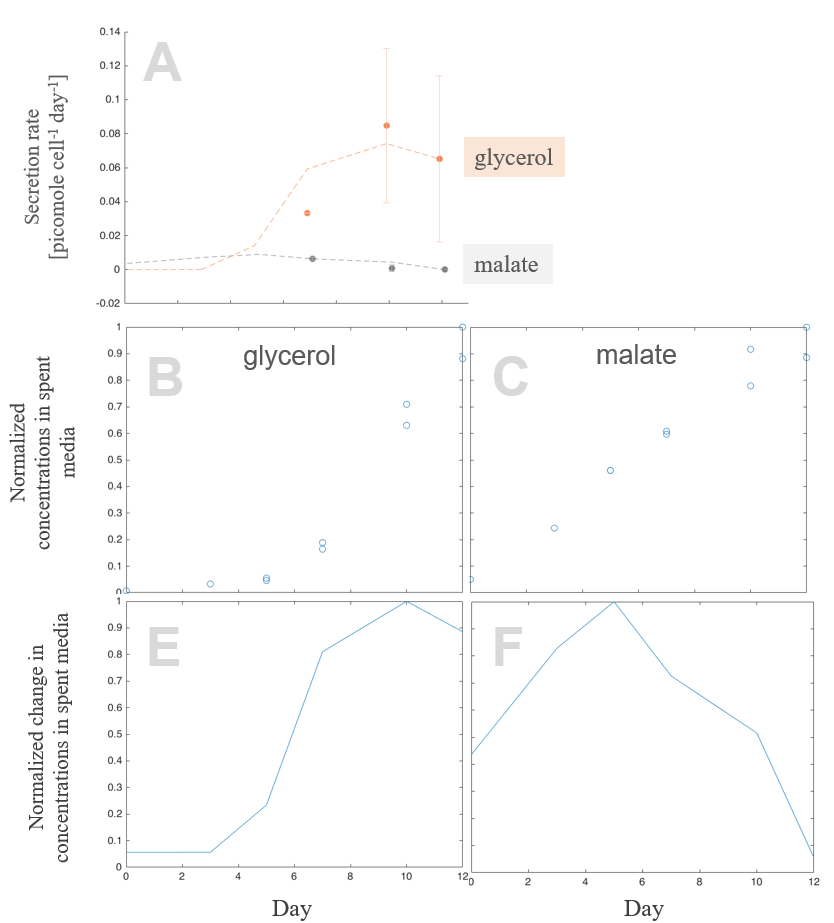

**Figure S11: Metabolomics data suggest glycerol secretion**

Accumulation of glycerol (*A*, red points), but not of malate (*A*, grey points), is observed in spent media from bioreactor runs for clones B1 and B2. The error bars represent the high and low values of metabolite consumption rates. Possible time-course profiles of consumption rates are also shown (*A*, dotted lines). These profiles were derived from consumption rate measurements from bioreactor runs for clones A1 and A2 (*E*, *F*), which were in turn calculated from concentration data quantified via mass spectrometry (*B*, *C*).

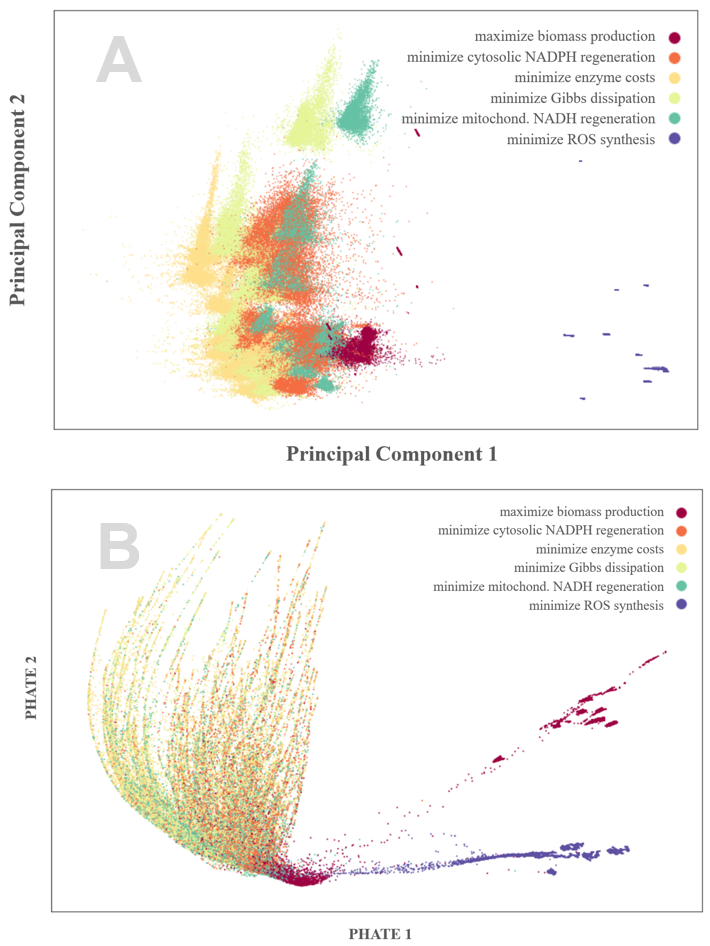

**Figure S12: Objective functions produce diverse intracellular flux distributions**

Intracellular flux distributions for 577 major reactions were compared using Principle Component Analysis (*A*) and PHATE (*B*). These visualizations show that *minimize ROS synthesis* (dark purple) produced intracellular flux distributions that diverged significantly from those of other objective functions. Intracellular fluxes of *maximize biomass production* (magenta) were dissimilar to a lesser degree from that of the remaining objective functions. Principle components 1 and 2 explained 4.2 and 3.3% of observed variance, respectively. PHATE visualization was obtained by setting the *knn* parameter to 100; the remaining parameters were left at default values.

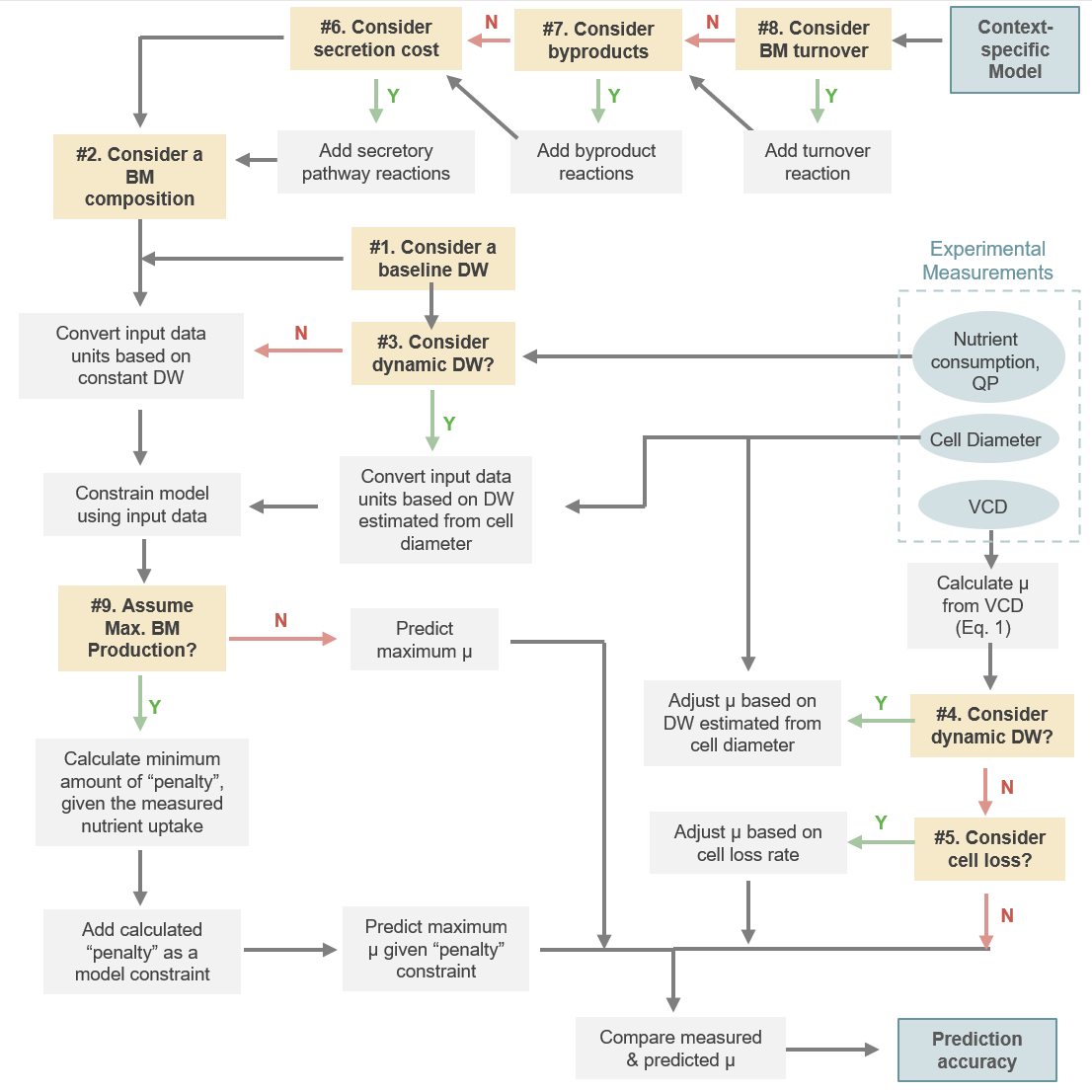

**Figure S13: Flowchart of Model Parameterization**

This figure describes the workflow and decision making behind model parameterization. It visualizes the *Model implementation of parameters* sub-section of the Methods section. The flowchart starts the context-specific model and experimental data in the (upper right), and ultimately ends by calculating prediction accuracy (lower left). The following abbreviations were used in the flowchart: DW, (cellular) dry weight; BM, biomass; μ, growth rate; VCD, viable cell density.
