## Supplementary material for "Systematic evaluation of parameters for genome-scale metabolic models of cultured mammalian cells": Annotated Pseudocode

Supplementary Documents: Annotated Codes for Flux Balance Analysis

### 1. Background

Irreversible model is required for alternative formulation of objective function

A metabolic network model describes cellular metabolism as a large-scale stoichiometric network of reactions and metabolites, and can be interrogated using constrain-based modeling techniques. A widely used constrain-based analysis approach is *flux balance analysis*, which formulates the model as a linear programming optimization problem by assuming an ‘objective function’. We found the conventional objective function of ‘biomass maximization’ to be inadequate in describing CHO metabolism, as discussed in the final report. There, we proposed an alternative objective function scheme that assumes biomass production is limited by cellular limitations such as enzyme capacity or ROS toxicity. This approach requires that reversible biochemical reactions to be described in a specific way, as elaborated below.

Metabolic network models can describe reversible biochemical reactions in two ways. For example, the second step of glycolysis interconverts glucose-6-phosphate and fructose-6-phosphate, catalyzed by glucose-6-phosphate isomerase. Notably, this is a reversible reaction, although the direction of fructose-6-phosphate production is favored in many cellular contexts. This reversible reaction, then, can be mathematically formulated either as a single bi-directional reaction (Table 1, eq. 1) or two unidirectional reactions (Table 1, eq. 2&3). Importantly, our modeling approach prerequisites an *irreversible* model that formulates reversible reactions as two unidirectional reactions. A metabolic network model can be converted to an irreversible model via the convertToIrreversible function from Cobra Toolbox^1^.

Table 1

| **#** | **Reaction name** | **Reaction formula** | **Description** |
| --- | --- | --- | --- |
| 1 | PGI | g6p[c] <-> f6p[c] + 52.992 kDa | Bi-directional reaction in a reversible model |
| 2 | PGI_f | g6p[c] -> f6p[c] + 52.992 kDa | Forward reaction in an irreversible model |
| 3 | PGI_b | f6p[c] -> g6p[c] + 52.992 kDa | Reverse reaction in an irreversible model |
| 4 | enzyme_cost | kDa -> | A ‘sink’ reaction that helps formulate ‘enzymatic cost’ as linear optimization problem |

A critical parameter of the model and flux balance analysis is the formulation of the objective function. We hypothesized that culture growth rate is bounded cellular limitations and have annotated the model accordingly. For example, the second step of glycolysis is annotated in the model by its ‘enzyme cost’, approximated by the mass of glucose-6-phosphate isomerase (Table 1, eq. 1-3; 52.992 kilodaltons).

The cumulative amount of ‘enzyme costs’ of all annotated metabolic reactions can be quantified and constrained via a ‘sink’ reaction (Table 1, eq. 4), a useful mathematical artifact that does not reflect any biological process. Through this ‘sink’ reaction, the total amount of ‘costs’ can be enumerated or manipulated, as previously demonstrated in parsimonious flux balance analysis and related methods^2^. This approach requires the model to be irreversible, as reversible reactions can erroneously create ‘negative enzyme cost’ by carrying flux in the reverse direction. In contrast, ‘enzyme cost’ can correctly be accounted for by formulating reversible reactions as two separate reactions. The implementations of other cellular limitations – such as ROS production or Gibbs energy dissipation – are analogous to this example.

### 2. Annotation

Setting up the analysis

An irreversible metabolic network model is loaded (9). The loaded model is a structure type variable with pertinent fields (rxns, mets, S etc.). Key variables are defined in preparation to the analysis. The cell array exceptRxns lists experimental data that should not be used as constraints (14). Nomenclature for the data are found earlier in the code (4-7). Here, we do not constrain growth rate ('mu') since we are predicting it. Additionally, we do not input cysteine consumption rate ('cys') due to it being difficult to measure accurately.

The variable metab_lim_name identifies the name of the sink reaction corresponding to an objective formulation (15). Sink reactions corresponding to other formulations include: 'gibbs_dissipation', 'ros_synthesis', 'cyto_nadph_synthesis', 'mito_nadh_synthesis', 'total_flux'. The variable contraint_margin describes the percentage margin with which experimentally measured input data will be overlayed with. For example, if the variable has a value of 15, glucose consumption rate of the model will be constrained to be within 15% of measured rates.

The variable min_norm is a parameter to the optimizeCbModel function (16). A default value of 0 leads to a conventional linear programming (LP) problem. Alternative values such as 'one' or 'zero' modify the LP problem; for more information, consult Cobra Toolbox documentation.

Experimental data of bioprocess measurements are loaded from CSV files via the loadExpmData function (20-25), which returns a table variable detailing the viable cell density, specific productivity, metabolite consumption rates, cell diameter and other variables. Crucially, the loadExpmData function converts measurement values to model-friendly units of mmol/gDW/hr. Lastly, a table variable is prepared to detail information regarding model predictions and results (27-33).

Model prediction of growth

The rest of the code loops through these data tables to make model predictions for each clone and timepoint of interest (36-100). Key steps in model prediction includes data input (60), application of limitation to proliferation (64-66), linear programming optimization (70-72) and saving prediction results in table form (75-96). In the case of modeling CPD17, an *in-silico* knockout of BCAT enzyme is realized by preventing isovalerate and 2-hydroxybutyrate to be secreted. This in turn, prevents the synthesis of these by-products, simulating the knock-out.

Data input onto model is realized by the constrain_irrev_model function according to the exceptRxns and constrain_margin variables defined above (60). The function adjusts values of the lower bound and upper bound fields of the model (lb and ub, respectively) according to measurement values. The function’s constrain_method parameter determines if the lower bound and/or upper bound constraints are applied. For example, the string 'cf' would constrain both lower bounds and upper bounds; the letters 'c' and 'f' stand for ceiling and floor respectively. The strings 'c' or 'f' by themselves would constrain only the lower bounds or upper bounds, respectively.

Given the inputted boundary conditions, a limit to proliferation is formulated by minimizing a previously determined cost value (64, 65). The resulting minimal cost value, found in the f field of the solution structure sol, is then set as an additional boundary condition (66). In the presented case, the minimum flux of enzyme_cost (Table 1, eq. 4) would be calculated and set as another boundary condition. Finally, given the inputted media substrate consumption rates and bounded enzyme capacity, a maximum growth rate is calculated (69-72). Importantly, the solved growth rate is adjusted to account for time-dependent changes in biomass (78) and viability (79-81), as discussed in the report. This adjusted growth rate, model with boundary conditions and resulting solution structure are all stored in a table to be outputted (75-96).

Adapting the presented code

The presented code predicts growth rate for inputted specific productivity and substrate consumption rates. The code can be easily altered to make similar predictions by changing which data is used as input and which bioprocess feature is predicted. This can be done by adjusting the the values of exceptRxns and obj_func. For example, exceptRxns and obj_func can be changed to {‘qp’} and {‘DM_igg[c]’} respectively to make predictions of specific productivity from inputted growth rate and substrate consumption rate. Alternatively, prediction of amino acid consumption rates was demonstrated by setting exceptRxns to amino acid consumption rates (excluding glutamate and glutamine) and setting metab_lim_name as ‘total_flux’. The realization of these adjustments can be found in the transferred code.

### 3. Code

```
function [Predicts, datas] = Predict_max_mu()
close all;
mets = {'asp','glu','cys','asn','ser','gln','his','gly','thr','arg','ala','tyr','val','met','trp','phe','ile','leu','lys','pro','glc','lac'};
abbrev = [{'mu', 'qp'}, mets];

%load model
load('pfizer_model.mat', 'pfizer_model');
metab_lim_name = 'enzyme_cost';
pfizer_model.metab_lim = zeros(size(pfizer_model.rxns));
pfizer_model.metab_lim(findRxnIDs(pfizer_model, metab_lim_name)) = 1;
exceptRxns = {'mu', 'cys'};
constrain_margin = 15;
min_norm = 0; % 'one';


%% ------ make predictions
%load data
data1 = loadExpmData('cpd5');
data2 = loadExpmData ('cpd9');
data3 = loadExpmData ('cpd11');
data4 = loadExpmData ('cpd17');
datas = {data1, data2, data3, data4};

%table
n = [1:80]';
data = zeros(size(n)); day = zeros(size(n)); replicate = zeros(size(n));
error = NaN(size(n));
model = cell(size(n)); sol = cell(size(n)); fluxtable = cell(size(n));
Predicts = table(n, data, day, replicate, model, sol, error);
c = 1;

days = [4 5 6 7 8 9 10 11];
for i = 1:length(datas)
    %treatment
    repl = unique(datas{i}.replicate);
    for k = 1:length(repl)
        %culture day
        for j = 1:length(days)
            %pull experimental data from table
            ix = intersect(find(datas{i}.day == days(j)) , find(datas{i}.replicate == repl(k)) );
            ix1 = intersect(find(datas{i}.day == (days(j)+1)) , find(datas{i}.replicate == repl(k)) );
            expm_mu = datas{i}.mu(ix);
            expm_qp = datas{i}.qp(ix);
            for a = 1:length(abbrev)
                Predicts{c, abbrev{a}} = datas{i}{ix,abbrev{a}};
            end
            Predicts.expm_mu(c) = expm_mu;
            Predicts.expm_qp(c) = expm_qp;
            Predicts.cellwt(c) = datas{i}.cellwt(ix);

            %make prediction
            model = pfizer_model;
            if i == 4
                model = changeRxnBounds(model, {'EX_2hb_e_', 'EX_iv_'}, 0, 'b');
            end
            model = constrain_irrev_model(model, datas{i} ,days(j), repl(k), exceptRxns, 'cf', constrain_margin);

            %apply metabolic limitation
            model = changeObjective(model, metab_lim_name);
            sol = optimizeCbModel(model, 'min');
            model = changeRxnBounds(model, metab_lim_name, sol.f, 'b');

            %solve model for prediction
            obj_func = 'biomass_cho_producing';
            model = changeObjective(model,  obj_func);
            sol = optimizeCbModel(model, 'max', min_norm);
            model = changeRxnBounds(model, obj_func, sol.f, 'b');

            %fill in table
            pred_mu = 0;
            if sol.stat == 1
                pred_mu = sol.f;
                pred_mu = pred_mu * datas{i}.cellwt(ix)/datas{i}.cellwt(ix1);
                if days(j) >= 6
                    pred_mu = pred_mu - 0.005;
                end
                pred_qp = sol.x(findRxnIDs(model, 'DM_igg[c]'));
                Predicts.pred_qp(c) = pred_qp;
                Predicts.cellwtfc(c) = datas{i}.cellwt(ix1)/datas{i}.cellwt(ix);
                Predicts.pred_mu(c) = pred_mu;
            else
                Predicts.pred_qp(c) = NaN;
            end


            %fill in table further
            Predicts.data(c) = i;
            Predicts.day(c) = days(j);
            Predicts.replicate(c) = repl(k);
            Predicts.model{c} = model;
            Predicts.sol{c} = sol;
            c = c+1;
        end
    end
end

SE = sum(abs(Predicts.error(~isnan(Predicts.error))));
SSE = sum(Predicts.error(~isnan(Predicts.error)).^2);
```
